# Gut Microbiota Production of Phenylacetate Programs Vascular Niche Senescence and Drives Atherosclerosis

**DOI:** 10.64898/2026.02.27.708541

**Authors:** Khatereh Shabanian, Florentin Constancias, Benoit Pugin, Taraneh Shabanian, Aurelien Thomas, Sylvain Le Gludic, Marianne Spalinger, Alessia Mongelli, Cristina Menni, Xinyuan Zhang, Lorenzo Da Dalt, Manuel Colucci, Hector Rodriguez Cetina Biefer, Omer Dzemali, Matthias Hermann, Andrea Alimonti, Giuseppe Danilo Norata, Francesco Paneni, Frank Ruschitzka, Seyed Soheil Saeedi Saravi

## Abstract

Vascular senescence is a key contributor to ageing-related diseases, including atherosclerosis. Initial intervention is based on aggressive management of traditional risk factors, yet microbial metabolites remain underestimated as modifiable factors. We recently identified phenylacetate (PAA), a gut microbiota–linked metabolite, as a potent accelerator of endothelial senescence, raising the question of its causal role in atherosclerosis. Here, we show that PAA promotes vascular niche senescence and perivascular adipose tissue (PVAT) dysfunction, associated with atherosclerosis in humans and mice. Furthermore, PAA administration to atherosclerosis-prone mice was sufficient to drive atherosclerosis without altering lipid profile. Mechanistically, we found that PAA induces senescence-messaging secretome, containing IL-6, from endothelial cells, which stimulates Notch1 and disrupts insulin signaling in adipocytes. Blocking the PAA–IL-6–Notch1 axis as well as senolytics rescued adipocyte senescence and dysfunction. Identification of the strong link between PAA and atherosclerosis opens new avenues for microbiome-targeted preventive and therapeutic strategies in ageing.

**Highlights:**

- Age-dependent increase in gut microbial metabolite PAA causally promotes vascular niche senescence
- PAA indirectly accelerates PVAT senescence through endothelial senescence-messaging secretome
- SASP upregulates Notch1 signaling, leading to PVAT dysfunction
- Senolytic therapy restores PVAT function
- PAA-induced vascular niche senescence contributes to atherosclerosis progression

## Introduction

Vascular dysfunction is thought to act as both early sensor and as driver of ageing throughout the body^1^. This underpins cardiovascular and metabolic diseases as leading causes of death in aging populations. Emerging evidence highlights the gut microbiome as a key modulator of vascular aging, with its composition and function shifting toward a pro-inflammatory and dysbiotic state over time^2,3^. These alterations in gut microbial ecology are associated with an increased systemic burden of microbial-derived metabolites, some of which exert profound driving effects on cardiovascular diseases (CVD), particularly atherosclerosis being its main determinant^4,5^. Among these, metabolites derived from aromatic amino acids (AAAs) have garnered significant attention for their roles in accelerating cardiovascular pathologies^2,6–8^. However, these metabolites and their precise mechanisms remain incompletely identified.

Among AAAs, phenylalanine-derived metabolites, particularly phenylacetic acid (PAA) and its downstream product phenylacetylglutamine (PAGln), have emerged as key regulators of cardiovascular risk in ageing^2,8^. We have recently reported that PAA elevates in aged humans and mice and causally induces aortic endothelial cell senescence^2^. Mechanistically, we demonstrated that PAA triggers mitochondrial oxidative stress, driving a senescence-associated secretory phenotype (SASP) that amplifies vascular dysfunction. These findings establish a direct link between gut microbiota-derived PAA and vascular aging, positioning this metabolite as a potential driver of atherosclerosis. Prior studies revealed that proteins secreted by aged vessels may accelerate cellular senescence throughout the other organs^9,10^. In line with this, we assume that PAA-induced endothelial senescence may potentially impact the adjacent cell types like adipocytes in perivascular adipose tissue (PVAT), triggering an inflammatory aortic microenvironment; yet, their crosstalk and mechanisms remain unknown. Given the critical role of PVAT in vascular inflammation and atherosclerosis^11–14^, understanding how PAA interacts with PVAT biology and function is critical for uncovering novel mechanisms and therapeutic targets for vascular aging and related atherosclerosis.

We, therefore, hypothesized that gut microbiota-derived PAA induces endothelial-PVAT senescence milieu, reinforcing atherosclerosis development. We sought to explore whether and how age-associated elevation of PAA disrupts PVAT homeostasis and function indirectly through endothelial cells, contributing to atherosclerosis progression. Our findings may identify PAA as a predictive biomarker for early stages of atherosclerosis and build a foundation for microbiome-targeted interventions aimed at preserving (peri)vascular function and mitigating atherosclerosis in aging populations.

## Results

### Gut microbiota alteration is associated with elevated PAA levels in ageing

We previously reported that gut microbiota-dependent phenylalanine metabolism and production of its metabolites increase with age^2^. Consistently, our untargeted LC-MS/MS metabolomics data revealed significantly elevated plasma concentrations of phenylalanine derivatives PAA (Fig. 1a) and PAGln (Extended Data Fig. 1a) in aged individuals enrolled in the TwinsUK aging cohort (*n*=2,953). Linear regression analysis showed that both metabolites are positively associated with age (PAA*: r* = 0.13, *R*^2^ = 0.017, *p*<0.001; PAGln: *r* = 0.27, *R*^2^ = 0.073, *p*<0.001). To validate the metabolic features of the ageing process observed in the TwinsUK study, we quantitated PAA concentration in plasma samples from a healthy validation cohort comprising 98 healthy individuals (Fig. 1b). Our targeted metabolomic data and correlation analysis confirmed a strong positive association between plasma PAA and age in this cohort (*R*^2^ = 0.4883, *p*<0.0001; Fig. 1c). Our data also showed a significantly higher plasma PAA concentrations in the aged population (>65□years old, *n*=45) compared to those in the young group (18-35□years old, *n*=41) (Fig. 1d). We next observed similar trends in data from our mouse chronological aging cohort. Old mice (24-30-months old) exhibited a significant elevation of plasma PAA levels compared to young mice (3-months old) (Fig. 1e). Notably, mice were matched for sex, body weight, and kidney function to rule out their effects on age-dependent PAA elevation in plasma, as reported previously^2^.

**Fig. 1.**
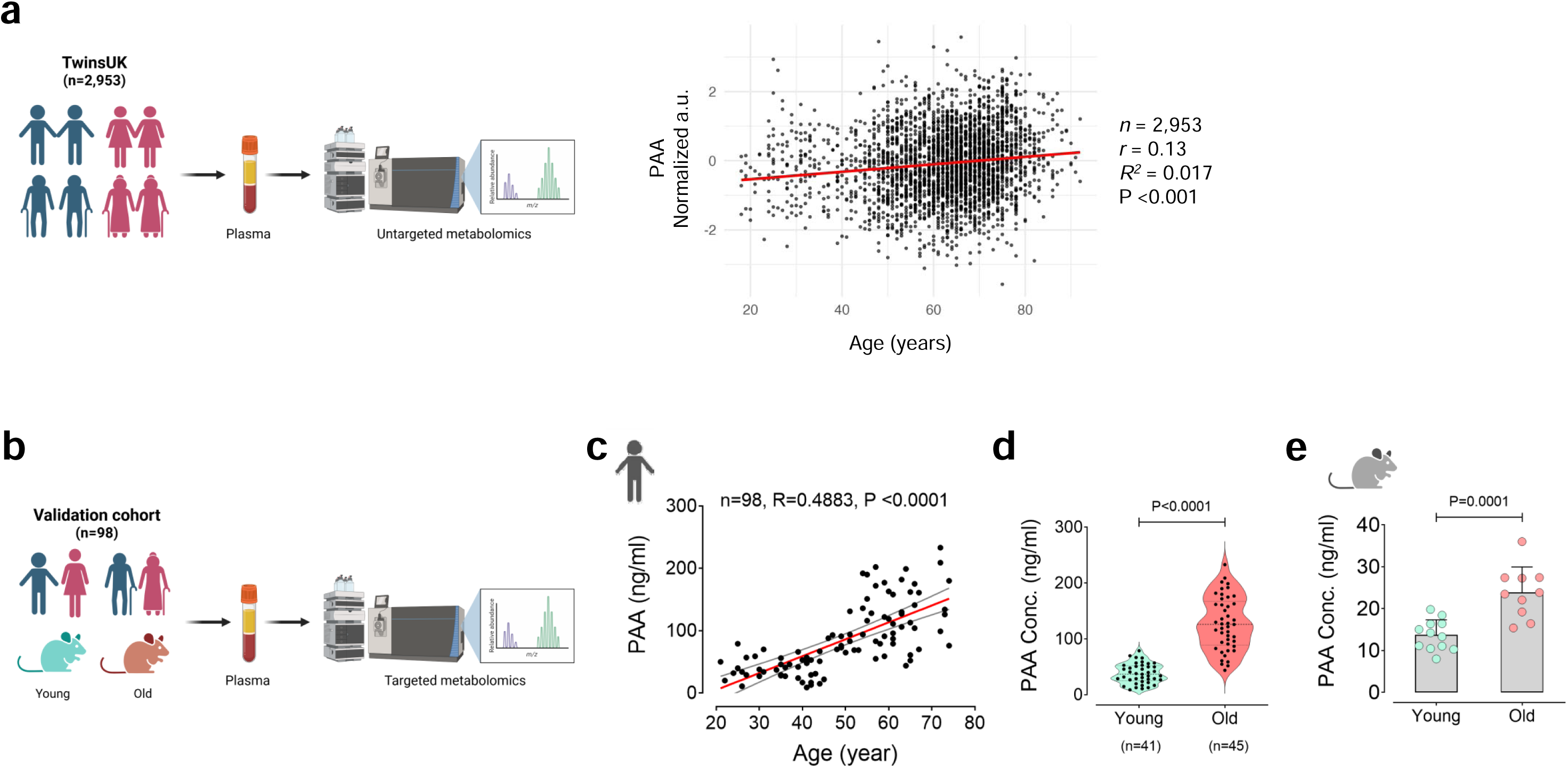
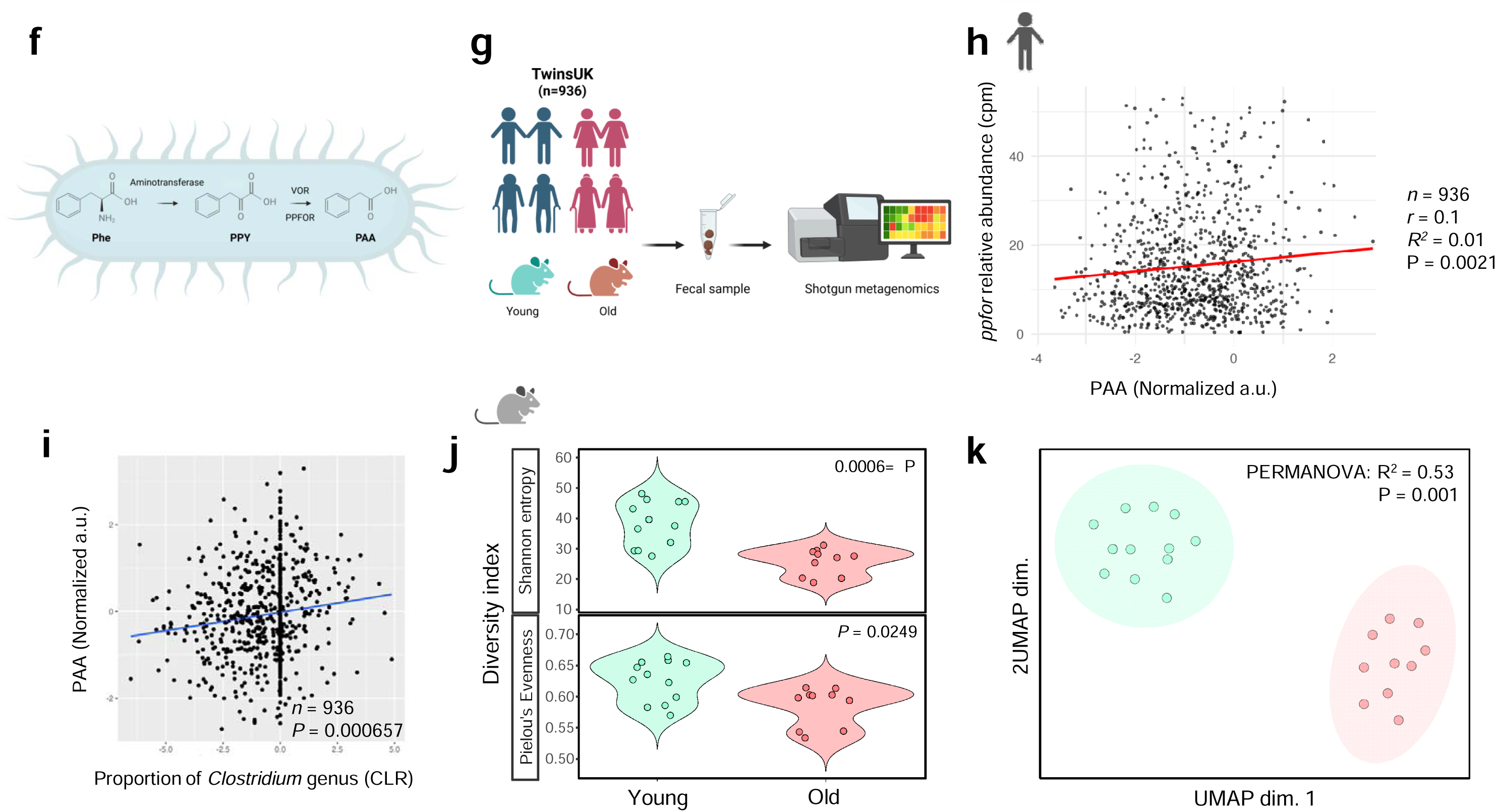
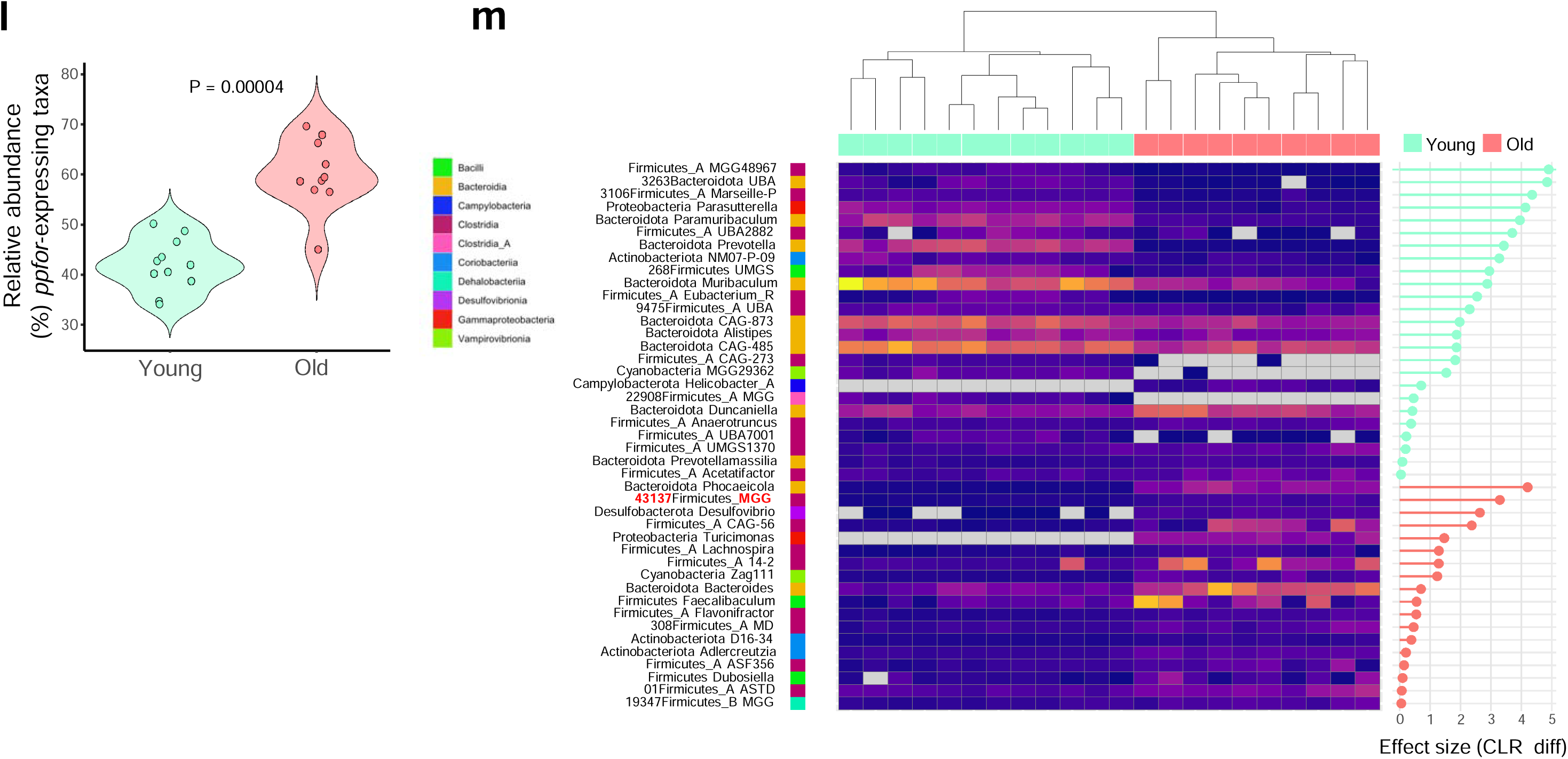
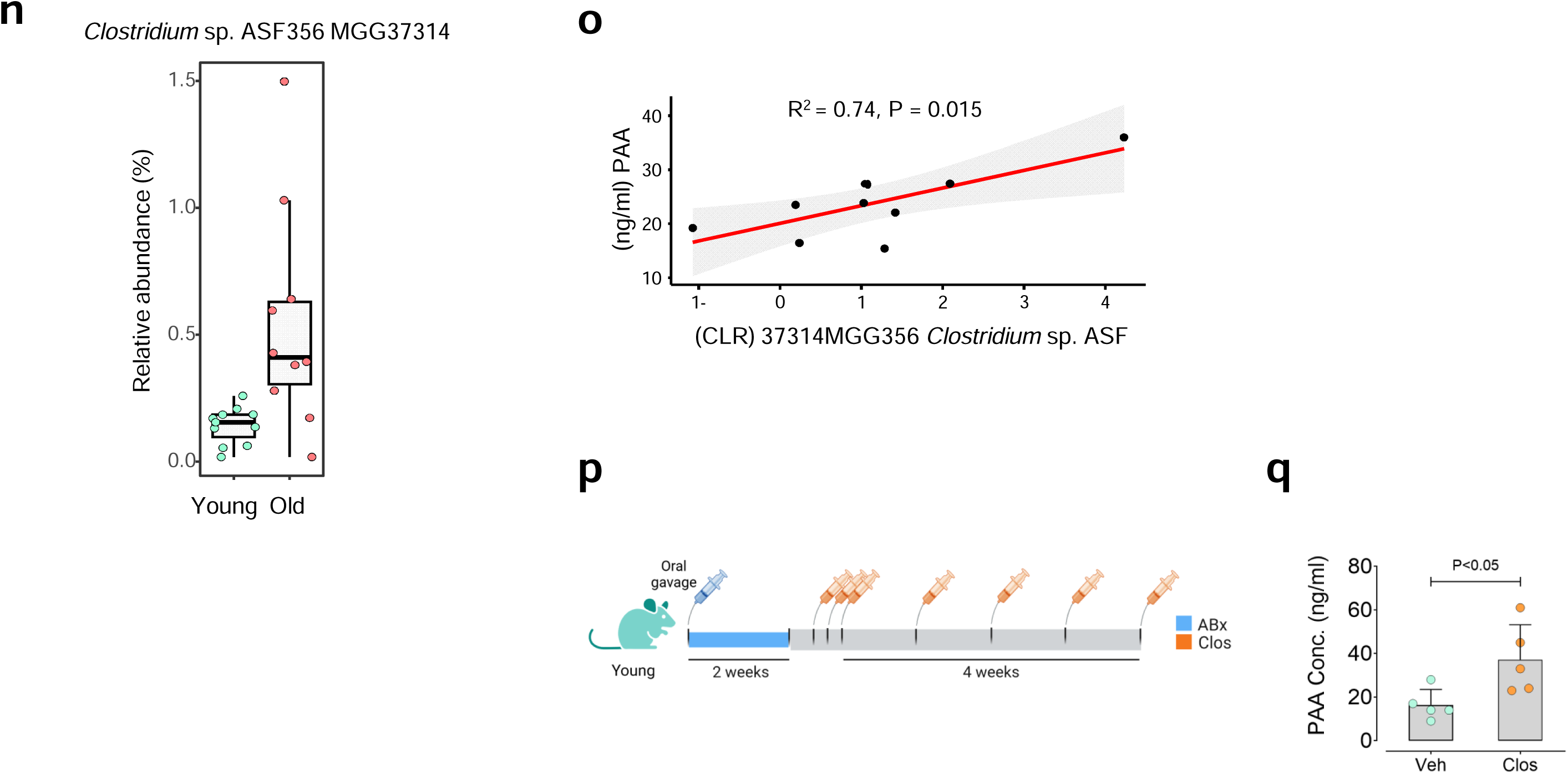
Aged gut microbial features is linked with plasma PAA elevation. **a,** Plasma samples from individuals enrolled in the TwinsUK aging cohort (*n* = 2,953; male and female) were subjected to untargeted metabolomics. Correlation between plasma PAA concentration and chronological age in the cohort. **b,** Plasma PAA concentration were quantified by targeted metabolome analysis in the healthy Validation cohort (*n* = 98; male and female) and in the mouse aging cohort (C57BL/6J male and female; *n* = 10-12). **c,** Correlation between the plasma PAA levels and chronological age in the Validation cohort (*n* = 98). **d,e,** Plasma PAA in healthy young (18-40 years) and old (>65 years) individuals from the Validation cohort (young, *n* = 41; old, *n* = 45) (**d**), and in young (3-month-old) and old (>27-month-old) mice (young, *n* = 12; old, *n* = 10) (**e**). **f,** Schema for key processes in conversion of dietary phenylalanine to PAA by VOR/PPFOR system in gut microbiota. **g,** Fecal samples were subjected to shotgun metagenomic sequencing in TwinsUK cohort (*n*□=□936) and mouse aging cohort (*n* = 10-12). **h,i,** Correlation between plasma PAA concentrations and relative abundance of *ppfor*^+^ bacteria (**h**) and abundance of *Clostridium* taxa (**i**) in TwinsUK cohort (*n*□=□936). **j,k,** Differences of alpha diversity measured by Shannon and Pielou’s evenness indices (**j**) and beta diversity based on the community level alterations (**k**) of gut microbiota among mouse aging cohort (*n*□=□10–12). **l,m,** Relative abundance (%) of *ppfor*^+^ taxa (**l**) and differential abundance of bacterial profiles (**m**) in the microbiomes of young (*n*□=□12) and old (*n*□=□10) mice. **n,** Relative abundance (%) of the *Clostridium* sp. ASF356 MGG37314 strain in the microbiomes of young (*n*□=□12) and old (*n*□=□10) mice. **o,** Correlation between plasma PAA concentrations and relative abundance of the *Clostridium* sp. ASF356 MGG37314 in mouse aging cohort (*n*□=□10–12). **p,** Young mice were pretreated with antibiotics (ABx) for 2 weeks, followed by colonization with *Clostridium* sp. ASF356 (*Clos*) for 4 weeks (*n*□=□5). **q,** Plasma PAA in *Clos*-mice (*n*□=□5) and vehicle-mice (*n*□=□5). Error bars represent SD (**e,n,q**), SEM (**j,k,l,n**), or 95% confidence intervals (**a,h,i**). Statistical analysis was performed using two-tailed Pearson correlation analysis (**a,h**), two-sided Spearman’s rank correlation test (**c,o**), two-tailed unpaired Student’s *t*-test (**d,e,l,n,q**), linear mixed model-centered log ratio (CLR) transformation (**i**), Kruskal–Wallis statistical test (**j**), Atchinson distance and permutational multivariate analysis of variance (PERMANOVA) (**k**), analysis of composition of microbiomes (ANCOM) method for microbial abundance analysis (**m**). Data are shown as median with min–max; each violin represent interquartile range (IQR); center lines indicate the median; violins extend from the min to max values (**d,j,l**). Images created with https://BioRender.com (**a,b,c,e-g,h,j,p**).

Our and others’ prior studies have reported that dietary phenylalanine (Phe) undergoes dehydration to produce phenylpyruvic acid (PPY) through the gut microbiota^2,7^. PPY is then converted into PAA mostly via two putative pathways involved in oxidative decarboxylation—phenylpyruvate:ferredoxin oxidoreductase (PPFOR) (KEGG orthology ID: K00179) and α-ketoisovalerate ferredoxin oxidoreductase (VOR) (KEGG orthology ID: K00169)^2,7,15^ (Fig. 1f). The direct contribution of gut bacteria to Phe metabolism has been previously confirmed by the antibiotic clearance of gut microbiota that led to suppressed PAA production in mice^2^. Furthermore, we conducted gene abundance annotation and taxonomic profiling of fecal shotgun metagenomic data in the TwinsUK cohort (*n*=936) (Fig. 1g). The *ppfor* gene contributing to catalytic conversion of PPY to PAA exhibited significant positive correlation with plasma PAA levels (*r* = 0.1, *R*^2^ = 0.01, *p*=0.0021; Fig. 1h). In light of these findings, we observed that 15 specific bacteria belonging to *ppfor*-expressing *Clostridium* genus (*Clostridium innocuum*, *Clostridium* sp. AF20_17LB, *Clostridium disporicum*, *Clostridium spiroforme*, *Clostridium* sp. AM33_3, *Clostridium* sp. AF27_2AA, *Clostridium SGB6179*, *Clostridium* sp. AF32_12BH, *Clostridium* sp. chh4_2, *Clostridium* sp. AF15_49, *Clostridium* sp. SN20*, Clostridium perfringens, Clostridium symbiosum,* and *Clostridium* sp. AM42_4) are enriched in older individuals (Fig. 1i, Extended Data Fig. 1b,c). Their strong positive correlation with plasma PAA concentrations suggests the capability of these bacteria to generate higher PAA levels with age (*p*=0.000657, Fig. 1i).

To deeply investigate the link between age-related functional alterations in the gut microbiota and the increases in circulating PAA, we conducted a metagenome analysis in our mouse model of ageing. Pronounced gut microbial feature shifts including alterations in both alpha-diversity (richness and evenness; Fig. 1j) and beta-diversity (Fig. 1k) indices were observed in old mice. Similar to what seen in our human cohort, we found an age-dependent increase in relative abundance (%) of *ppfor* gene homologs in the microbiomes (Fig. 1l). A centered log-ratio (CLR) by analysis of composition of microbiomes (ANCOM) identified 6 bacterial species from the *Clostridium* genus enriched in old group (Fig. 1m; Extended Data Fig. 2). Among these species, only one species, *Clostridium* sp. ASF356 MGG37314 (*Clos*), was moderately abundant in old microbiomes (mean: 0.07918%; Fig. 1n), and exhibited the strongest positive correlation with plasma PAA levels (*R^2^*= 0.74, *p*=0.015; Fig. 1o). Our previous work also validated a strong functionality of this species to convert L-Phenylalanine into PAA in BHI or YFCA culture media under anaerobic condition^2^. As a proof of concept, we pre-treated young mice with a broad-spectrum antibiotics cocktail (ABx) for 2 weeks, followed by weekly oral gavage of either *Clos* (5□×□10^8^) or vehicle for 4 weeks (Fig. 1p). The colonization led to significant elevation of plasma PAA (∼2.27-fold) in these mice (Fig. 1q), confirming metabolic functionality of the bacterium in converting dietary Phe to PAA *in vivo*.

These findings collectively suggest that the *ppfor*-expressing bacteria involved in Phe → PAA conversion enrich in the gut microbiota of aged humans and mice. Among these, *Clostridium* sp. ASF356 exhibits a strong ability to produce PAA in aged organisms.

### *Clostridium* sp. ASF356 through PAA induces premature PVAT senescence

Our data revealed that *Clos* and its metabolite PAA causally recapitulated the key features of PVAT senescence phenotype observed in aged mice (Extended Data Fig. 3a-c). We pre-treated young mice with ABx, followed by *Clos* colonization for 4 weeks (Fig. 2a). Upon tissue harvesting, we noted an increased vascularity of aortic PVAT. This observation was corroborated by senescence phenotype, evidenced by significant upregulation of (1) cell-cycle arrest markers CDKN2A (p16^INK4a^), CDKN2D (p19^INK4d^), and CDKN1A (p21^WAF1/Cip1^) at both mRNA and protein (for p16^INK4a^) levels (Fig. 2b,c), and (2) the SASP components, particularly IL-1α, IL-1β, IL-6, and CCL2, at protein level (Fig. 2d).

**Fig. 2.**
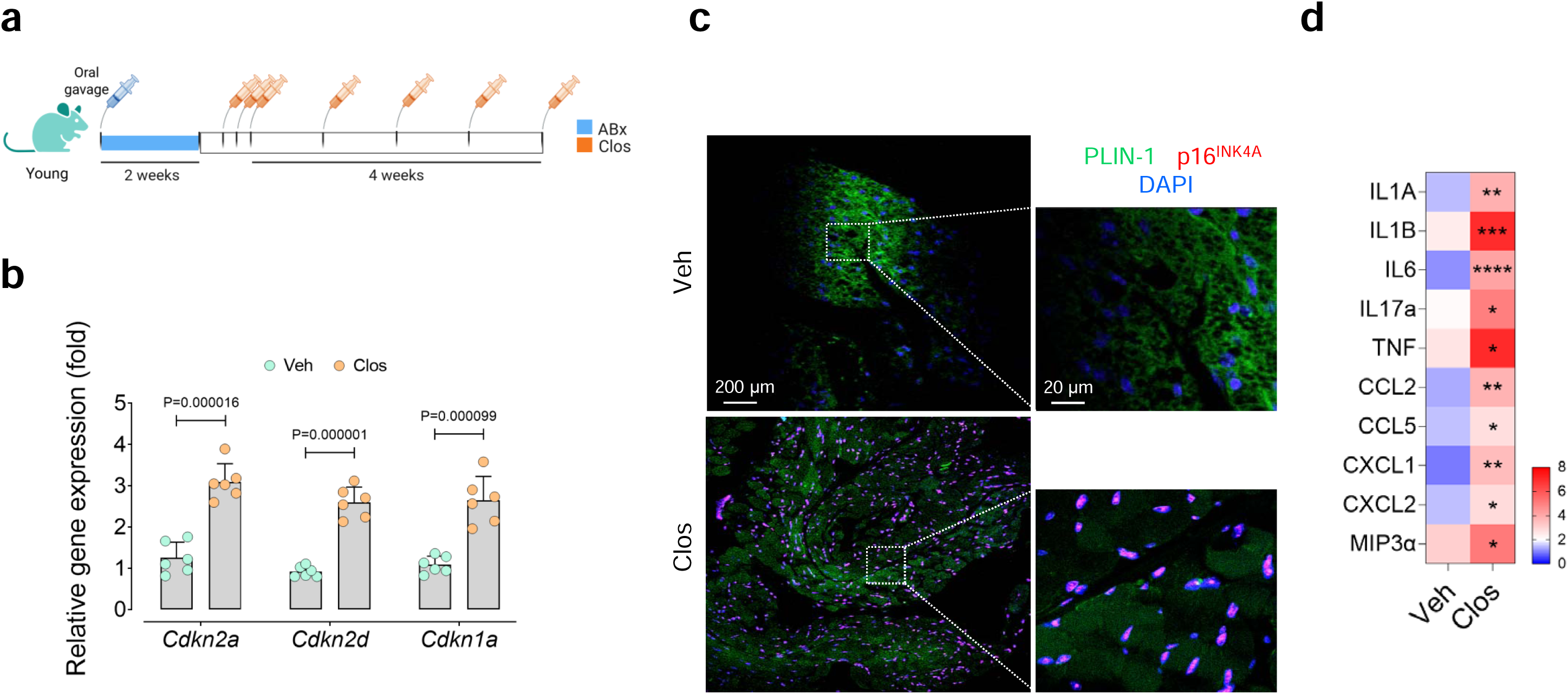
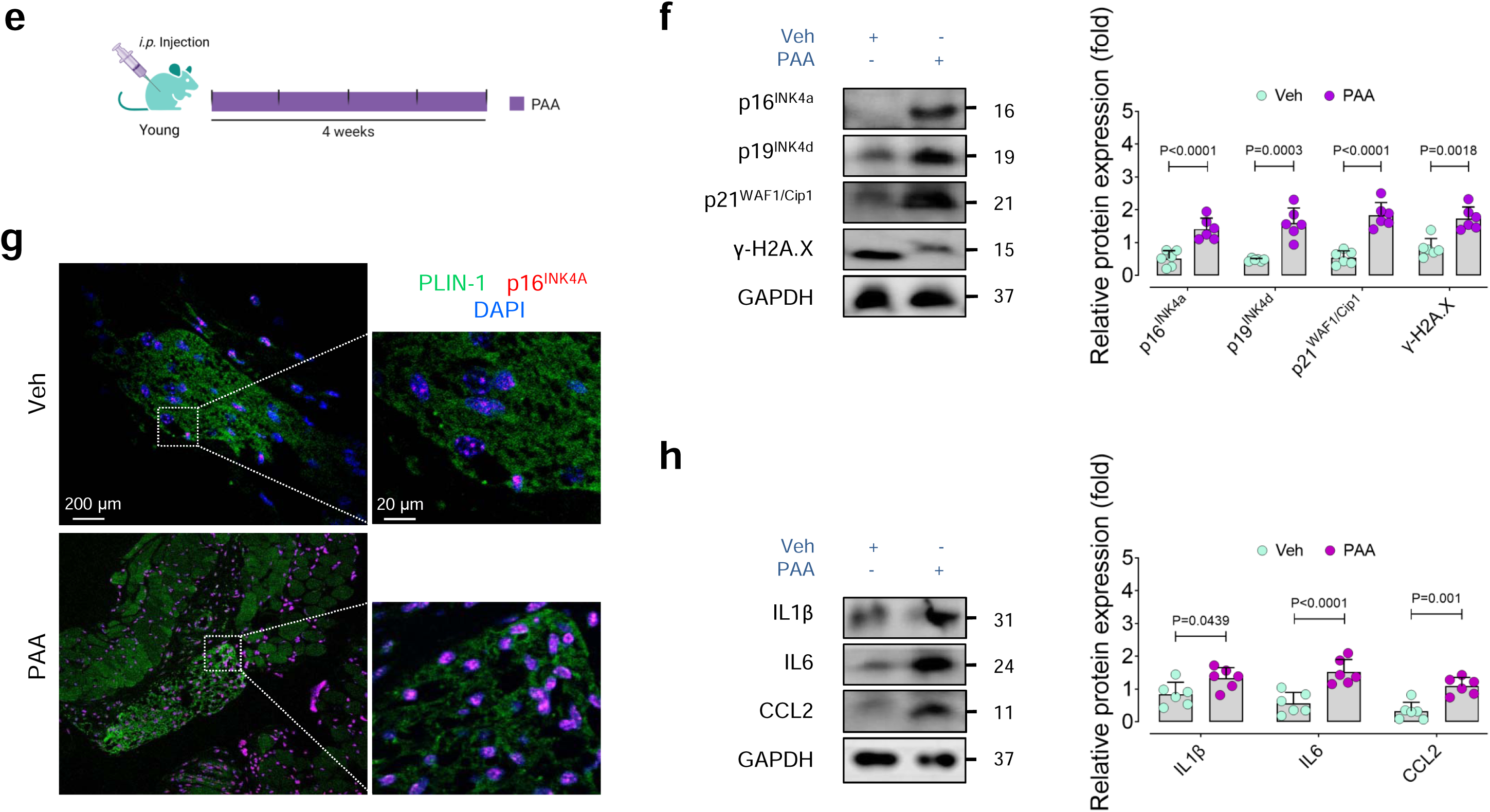
*Clostridium* sp. ASF356, through PAA, triggers aortic PVAT senescence. **a,** Young mice were pretreated with ABx for 2 weeks, and then colonized with *Clos* for 4 weeks (*n*□=□6). **b,** qPCR shows transcriptional alterations of CDK inhibitors *Cdkn2a*, *Cdkn2d*, and *Cdkn1a* in tPVAT from *Clos*-mice (*n* = 6) and vehicle-mice (*n* = 6). **c,** Representative confocal immunofluorescence images of p16^INK4A^ in tPVAT from these mice (*n* = 5). **d,** Relative protein expression analysis of SASP components in tPVAT from *Clos*-mice (*n* = 5) and vehicle-mice (*n* = 5). **e,** Young mice were administered to PAA (50 mg/kg, *i.p.*) daily for 4 weeks (*n*□=□6). **f,** Representative immunoblots and quantification of intensities for CDK inhibitors p16^INK4A^, p19^INK4D^, and p21^WAF1/Cip1^ and DNA damage marker γ-H2A.X in tPVAT from these mice (*n* = 6). **g,** Representative confocal images of p16^INK4A^ in tPVAT from these mice (*n* = 6). **h,** Immunoblotting represents the expression of the SASP components IL-1β, IL-6, and CCL2 in tPVAT from PAA-or vehicle-treated mice (*n* = 6). Data were determined in 8-10 micrographs and represent triplicated biologically independent experiments (**c,g**). Scale bars, 20 and 200 μm (**c,g**). Error bars represent SD (**b,d,f,h**). *P* values were calculated using a two-tailed unpaired Student’s *t*-test (**b,d,f,h**). Images created with https://BioRender.com (**a,e**). (\**P*<0.05, \*\**P*<0.01, \*\*\**P*<0.001, \*\*\*\**P*<0.0001).

To confirm that PAA is the underlying contributor of *Clos* to induce PVAT senescence, we treated young mice with PAA for 4 weeks (Fig. 2e). Our findings revealed that PAA confers senescence-promoting effects in thoracic PVAT (tPVAT) *in vivo*. Accordingly, PAA administration led to significant increase in the expression of cell-cycle arrest markers p16^INK4a^, p19^INK4d^, and p21^WAF1/Cip1^, alongside DNA damage response (DDR) marker γ-H2A.X (Fig. 2f). Cell-cycle arrest was also confirmed by p16^INK4a^ immunostaining (Fig. 2g), mirroring observations in tPVAT from *Clos*-colonized mice (Fig. 2c). Furthermore, PAA-treated mice exhibited a significant increase in the expression of the SASP components IL-1β, IL-6, and CCL2 in their tPVAT, similar to that observed in *Clos*-colonized mice (Fig. 2h).

Together, these data provide strong support for a causal contribution of *Clostridium* sp. ASF356 to aortic PVAT senescence *in vivo*, through enhanced phenylalanine → PAA conversion.

### PAA induces PVAT dysfunction by triggering endothelial senescence

Prior investigations revealed that age-associated endothelial cell (EC) senescence can directly trigger the adipocyte senescence and dysfunction through its senescence-messaging secretome^9,10^. Additionally, our previous study demonstrated that age-related increase in circulating PAA levels causes aortic EC senescence^2^. We, therefore, hypothesized that PAA-induced EC senescence might induce PVAT dysfunction through senescence-promoting activity. To examine this, we stimulated proliferating human aortic ECs (PEC; p.4-5) with exogenous PAA to induce premature senescence. As shown previously^2^ and in Extended Data Fig. 4a-e, PAA-exposed ECs exhibited the senescence signature phenotype including morphological changes (enlarged, flattened, increased granularity, and multinucleated appearance) and exacerbated senescence parameters comprising lysosomal alterations, reduced proliferation, and the SASP. When treated with premature senescent EC-conditioned medium (PAA-CM), human adipocyte-derived stem cell (hADSC)-derived adipocytes exhibited multiple senescence phenotype features (Fig. 3a), including reduced numbers of Ki67^+^ cells (Fig. 3b), increased SA-β-gal activity (32.21% *vs.* 6.11%, control-CM; Fig. 3c,d), and upregulated cyclin-dependent kinase (CDK) inhibitors *Cdkn2a*, *Cdkn2d*, and *Cdkn1a* (Fig. 3e). Senescence of adipocytes was also validated through increased DNA damage, evidenced by higher γ-H2A.X phosphorylation compared with cells treated with culture medium derived from vehicle-exposed PECs (control-CM; Fig. 3f). To further support these findings that CM derived from PAA-exposed ECs accelerates adipocyte senescence, we tested whether senolytics intervention with Dasatinib+Quercetin can selectively eliminate senescent adipocytes triggered by PAA-CM (Extended Data Fig. 5a). Our results demonstrated that D+Q selectively reduced viability of senescent ECs (∼64.89%), while sparing proliferating cells (∼85.96%) (Extended Data Fig. 5b).

**Fig. 3.**
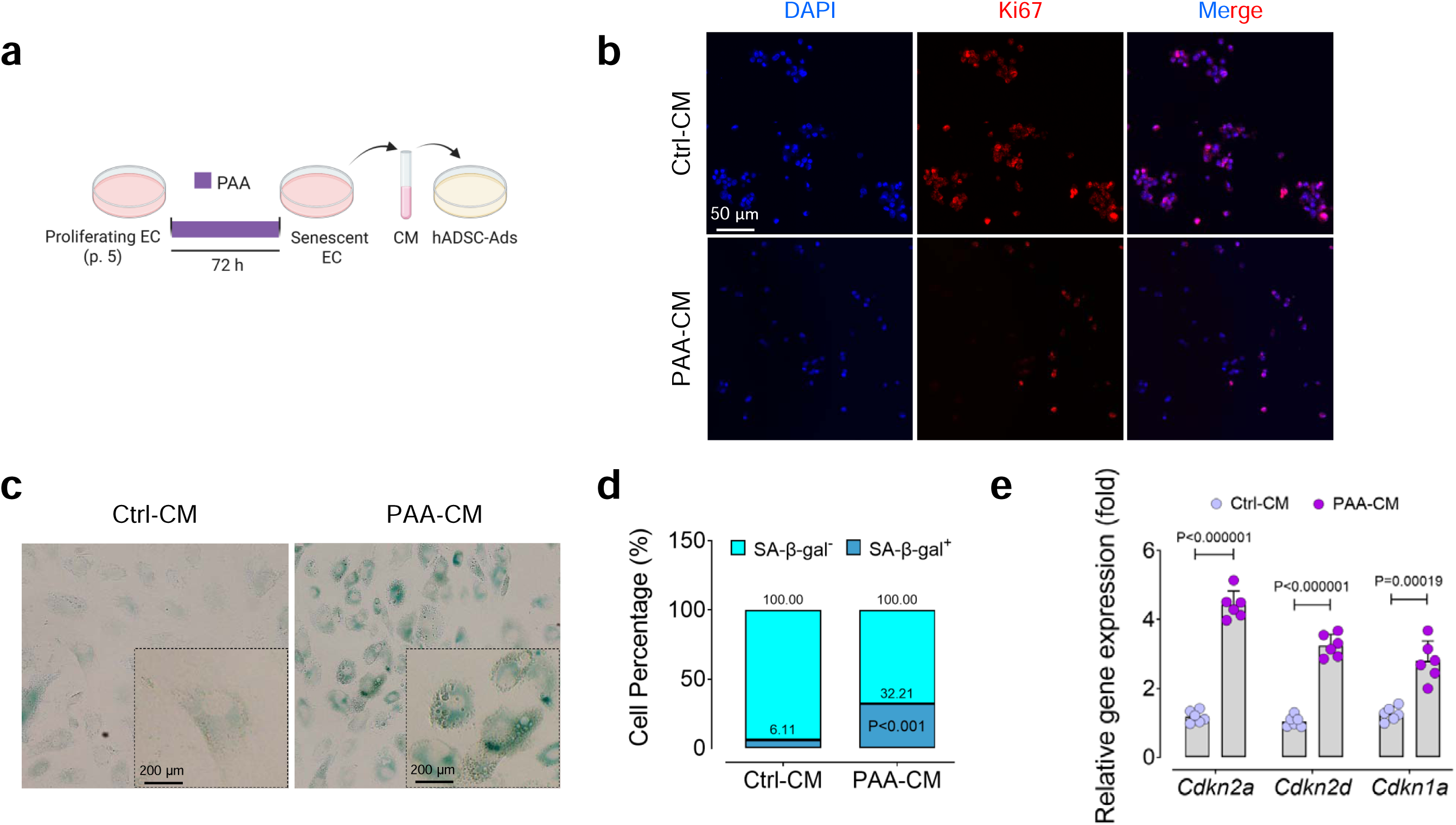
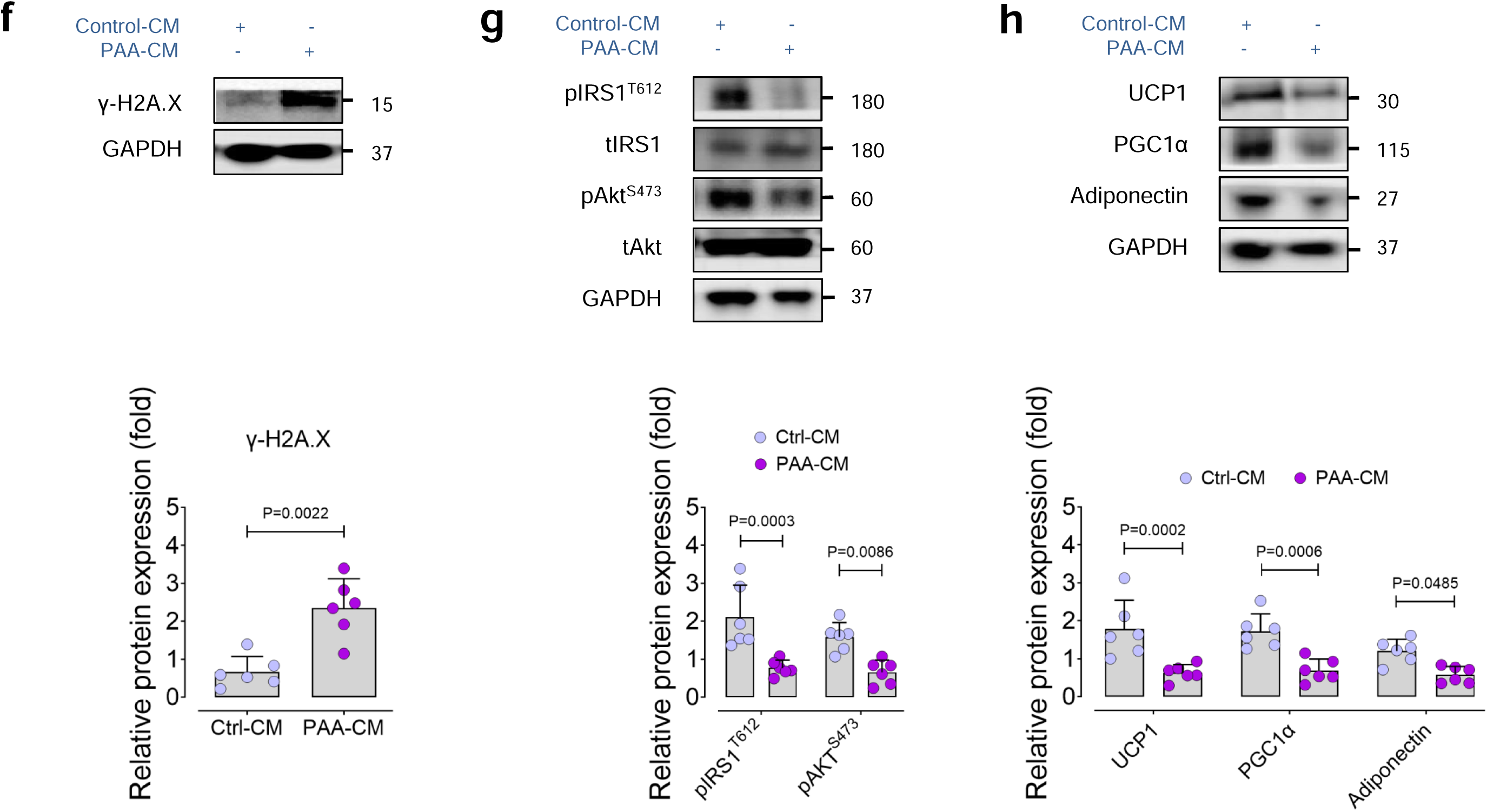
PAA indirectly induces adipocyte senescence and dysfunction. **a,** hADSC-derived adipocytes were treated with the conditioned medium (CM) derived from ECs (passages 4-5) exposed to PAA (10 μM, PAA-CM) or vehicle (control-CM) for 72 h. Cells were then subjected to senescence hallmarks profiling. **b,** Immunostaining reveals the expression of the proliferation marker Ki67 (*n* = 6). **c,** Representative bright-field images of SA-β-gal^+^ adipocytes treated with PAA-CM or control-CM, confirming senescence-associated lysosomal alterations. **d,** Quantitative plot is shown for percentage (%) of SA-β-gal^+^ *vs.* SA-β-gal^-^ cells (*n* = 6). **e,** qPCR demonstrates transcriptional changes of CDK inhibitors *Cdkn2a*, *Cdkn2d*, and *Cdkn1a* (*n* = 6). **f-h,** Representative immunoblots and quantification of intensities for DDR marker γ-H2A.X (**f**), the insulin signaling pathway (in the presence of insulin) (**g**), and proteins associated with adipocyte thermogenic function, including UCP1 and PGC1α (**h**) in adipocytes treated with the control medium or CM derived from PAA-exposed EC (*n* = 6). Data were determined in 8-10 micrographs and represent triplicated biologically independent experiments. Scale bar, 50 and 200 μm (**b,c**). Error bars represent SD (**d-h**). *P* values were calculated using a two-tailed unpaired Student’s *t*-test (**d-h**). Images created with https://BioRender.com (**a**).

As a proof of concept for senescence-promoting effects of PAA-CM, our *in vitro* studies revealed that hADSC-preadipocytes exposed to PAA-CM mimic senescence phenotype triggered by CM from H_2_O_2_-treated ECs (H_2_O_2_-CM; as a positive control to induce premature senescence) (Extended Data Fig. 6a). Notably, PAA triggers EC senescence by stimulating NOX4-mediated H_2_O_2_ generation, which were reported previously^2^. Our qPCR analysis confirmed that PAA-CM significantly increases transcriptional levels of the CDK inhibitors *Cdkn2a*, *Cdkn2d*, and *Cdkn1a* (Extended Data Fig. 6b), as well as the SASP components *IL1*Α, *IL1*Β, and *IL6* (Extended Data Fig. 6c) to the magnitude seen in response to H_2_O_2_-CM. Furthermore, preadipocytes treated with PAA-CM displayed markedly distinct senescence-like morphology and increased percentage of SA-β-gal activity comparable to those in H_2_O_2_-CM-treated cells (Extended Data Fig. 6d).

Importantly, hADSC-adipocytes in senescence-like state exhibited impaired cellular function including insulin signaling in association with reduced phosphorylation of insulin receptor substrate-1 (IRS1) at Tyr612 and its downstream protein kinase B (Akt) at Ser473 (Fig. 3g). Significant suppression was also observed in the expression of the browning and non-shivering thermogenesis marker UCP1 (uncoupling protein 1), alongside blunted adipogenic and thermogenic marker PGC1α (peroxisome proliferator-activated receptor gamma coactivator 1-alpha) and vasoprotective marker adiponectin in PAA-CM-exposed adipocytes (Fig. 3h). In contrast, treatment with control-CM did not induce such dysfunction features in adipocytes.

### Endothelial secretome triggers PVAT dysfunction through Notch1 signaling

To deeply understand the mechanisms by which senescent ECs trigger adipocyte senescence and dysfunction, we analyzed PAA-CM and found that the medium is enriched with senescence-messaging secretome. Our data revealed that IL-6 was abundantly secreted into the CM of senescent ECs stimulated by PAA, to the magnitude measured in replicative senescent ECs (p.15-17) (Fig. 4a). These findings are consistent with our previous results demonstrating that PAA upregulates IL-6 through HDAC4-mediated epigenetic modulation in ECs^2^.

**Fig. 4.**
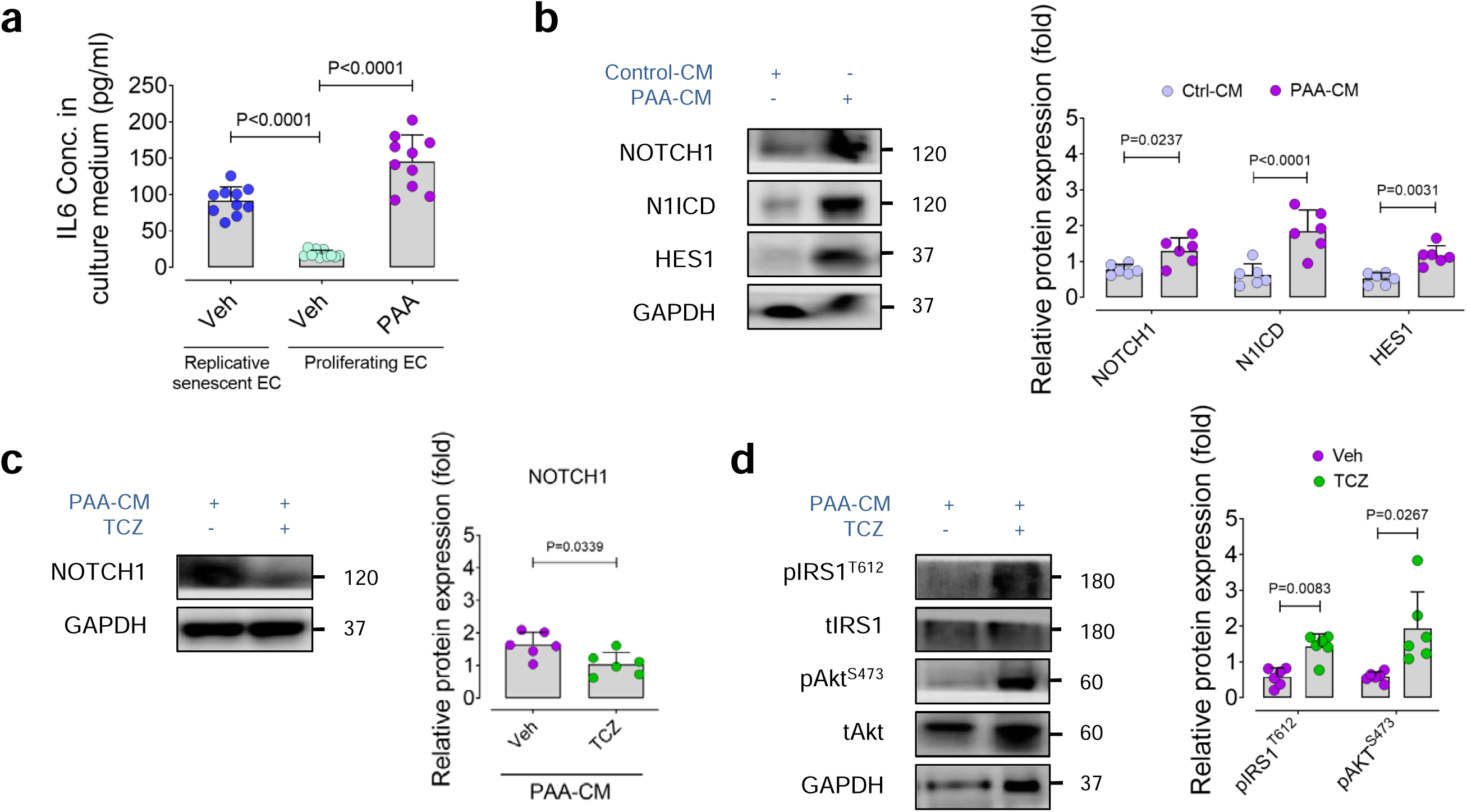
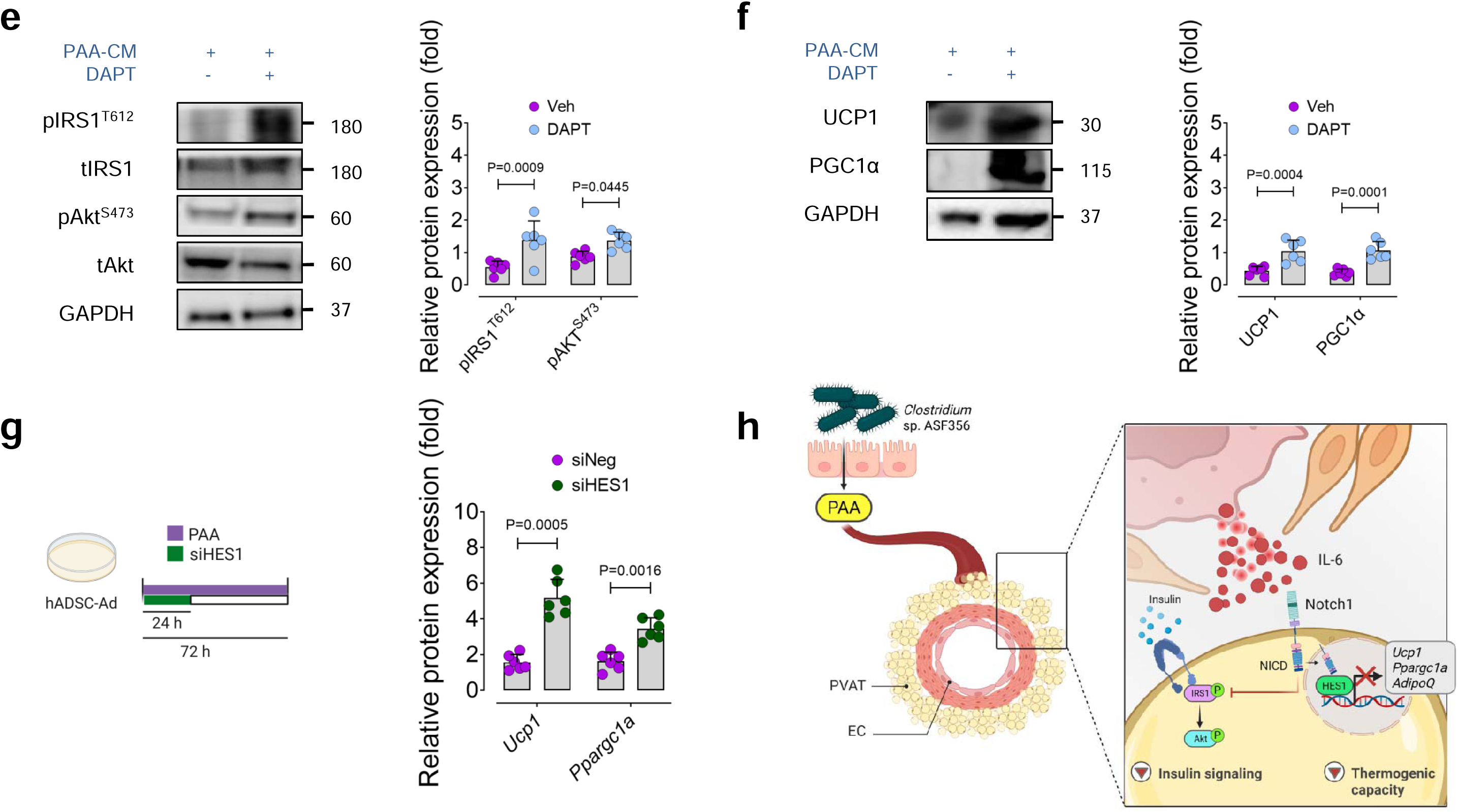
Premature senescent EC disrupts adipocyte function through the SASP-Notch signaling. **a,** IL-6 concentration in culture medium derived from replicative senescent or proliferating ECs treated with PAA (10 μM) or vehicle for 72 h (*n*□=□10 biologically independent samples). **b,** Representative immunoblots demonstrate the expression of NOTCH1 and downstream targets N1ICD and HES1 in hADSC-adipocytes exposed to CM derived from vehicle- or PAA-treated ECs (*n* = 6). **c,d,** Immunoblotting for NOTCH1 (**c**) and the insulin signaling pathway (**d**) in insulin-stimulated adipocytes treated with PAA-CM in the presence or absence of anti-IL6R neutralizing antibody, Tocilizumab (100 μg/mL) (*n* = 6). **e,f,** Immunoblotting for the insulin signaling pathway (**e**) and thermogenic markers UCP1 and PGC1α (**f**) in adipocytes treated with PAA-CM in the presence or absence of Notch inhibitor, DAPT (10 µM) (*n* = 6). **g,** qPCR represents transcriptional changes of thermogenic markers *Ucp1* and *Ppargc1a* in PAA-CM-exposed adipocytes transfected with siHES1 or siNeg (*n* = 6). **h,** Summary scheme outlining the mechanisms of microbial metabolite PAA for triggering adipocyte dysfunction. PAA indirectly activates NOTCH1 and its downstream HES1 in adipocytes by releasing senescence-messaging secretome containing IL6 from adjacent ECs. This both downregulates thermogenic function and the insulin signaling pathway in adipocytes. Error bars represent SD (**a-g**). *P* values were calculated using one-way ANOVA followed by Tukey’s post *hoc* test (**a**) and a two-tailed unpaired Student’s *t*-test (**b-g**). Image created with https://BioRender.com (**h**).

Ageing and obesity are both associated with a profound remodeling of PVAT secretome from vasoprotective profile, including e.g. adiponectin and nitric oxide, to a proatherogenic milieu enriched in IL-6 and resistin, thereby driving vascular decline^16^. Yet, the molecular mechanisms underlying age-associated PVAT dysfunction remain poorly understood. Recent evidence links IL-6 to Notch-mediated adipocyte dysfunction, disrupting adipogenesis and insulin sensitivity^17–19^. Accordingly, we tested whether and how IL-6 released from PAA-triggered senescent ECs promotes PVAT dysfunction by impairing insulin signaling and thermogenic capacity. Our results support the hypothesis that PAA-CM containing high IL-6 causes the upregulation of NOTCH1 signaling in hADSC-adipocytes, represented by increased expression of NOTCH1, Notch1 intracellular domain (N1ICD), and the downstream effector, hairy and enhancer of split 1 (HES1) (Fig. 4b).

To confirm that PAA-CM activates NOTCH1 signaling in an IL-6-depedent fashion, we examined PAA-CM-exposed adipocytes in the presence or absence of anti-IL6R neutralizing antibody, Tocilizumab (TCZ). Pharmacological IL-6 receptor inhibition markedly abolished PAA-mediated NOTCH1 activation, confirming that IL-6 is required for this effect (Fig. 4c). As a proof of concept, human recombinant IL-6 (hrIL6) recapitulated the PAA-induced activation of NOTCH1 in these cells (Extended Data Fig. 7a,b). We also tend to investigate how PAA regulates insulin signaling and thermogenic function. As shown in Fig. 4d, co-treatment with TCZ restored insulin signaling in PAA-CM-exposed adipocytes, as evidenced by recovered IRS1^T612^ and Akt^S473^ phosphorylation. In line with this, inhibition of NOTCH1 by DAPT—a pharmacological inhibitor of both NOTCH1 and γ-secretase, the protease complex responsible for releasing N1ICD^16^—abrogated the progressive suppression of insulin signaling and key thermogenic markers (mitochondrial UCP1 and PGC1α) triggered by PAA-CM (Fig. 4e,f). These were corroborated by reduced proportion of senescent cells in hrIL6+DAPT-*versus* hrIL6-treated cultures (SA-β-gal^+^ cells: 24.23% *vs.* 15.47%) and preserved insulin signaling (Extended Data Fig. 7c,d). To further explore the mechanistic link between NOTCH1 activation and impaired thermogenic programming, we silenced HES1 in PAA-CM–exposed adipocytes. HES1 knockdown significantly restored transcription of *Ucp1* and *Ppargc1a* compared to siNeg controls, demonstrating that PAA suppresses thermogenic gene expression through the IL-6–NOTCH1–HES1 axis (Fig. 4g).

These findings collectively suggest that PAA by elevating IL-6 acts as a critical paracrine mediator of endothelial-to-adipocyte signaling, promoting NOTCH1-HES1 signaling and thereby impairing insulin signaling and thermogenic programming in adipocytes (Fig. 4h).

### Senolytics rescue PAA-induced PVAT dysfunction

To both establish causality between *Clos* colonization and PVAT senescence and to explore a therapeutic avenue for reducing senescent cell burden, we exploited the potent senolytic D+Q cocktail in *Clos*-colonized mice (Fig. 5a). Our data revealed that D+Q treatment robustly attenuated key senescence hallmarks in tPVAT, including significant reductions in the expression of CDK inhibitors p16^INK4a^, p19^INK4d^, and p21^WAF1/Cip1^, as well as the phosphorylation of the DDR marker γ-H2A.X (Fig. 5b). This senescence rescue was accompanied by marked suppression of the SASP, as evidenced by reduced IL-6 signals (Fig. 5c), together with a pronounced decrease in lysosomal SA-β-gal activity in both aorta and surrounding PVAT (Fig. 5d). Importantly, our immunostaining analyses confirms that the elimination of senescent cells is associated with downregulation of NOTCH1 signaling in perivascular adipocytes (Fig. 5e). As earlier shown in Fig. 4, NOTCH1 acts as a key regulator of adipocyte insulin signaling and thermogenic pathway. We therefore hypothesized that D+Q-mediated suppression of NOTCH1 may restore PVAT metabolic function in *Clos*-colonized mice. Consistent with this notion and with our previous observations in vascular ECs^2^, senolytic clearance of *Clos*-triggered senescent cells led to a marked recovery of adipocyte function. Both insulin signaling and thermogenic capacity were significantly improved, as indicated by enhanced IRS1^T612^—Akt^S473^ phosphorylation and upregulation of UCP1 and PGC1α following 4 weeks of D+Q administration (Fig. 5f,g).

**Fig. 5.**
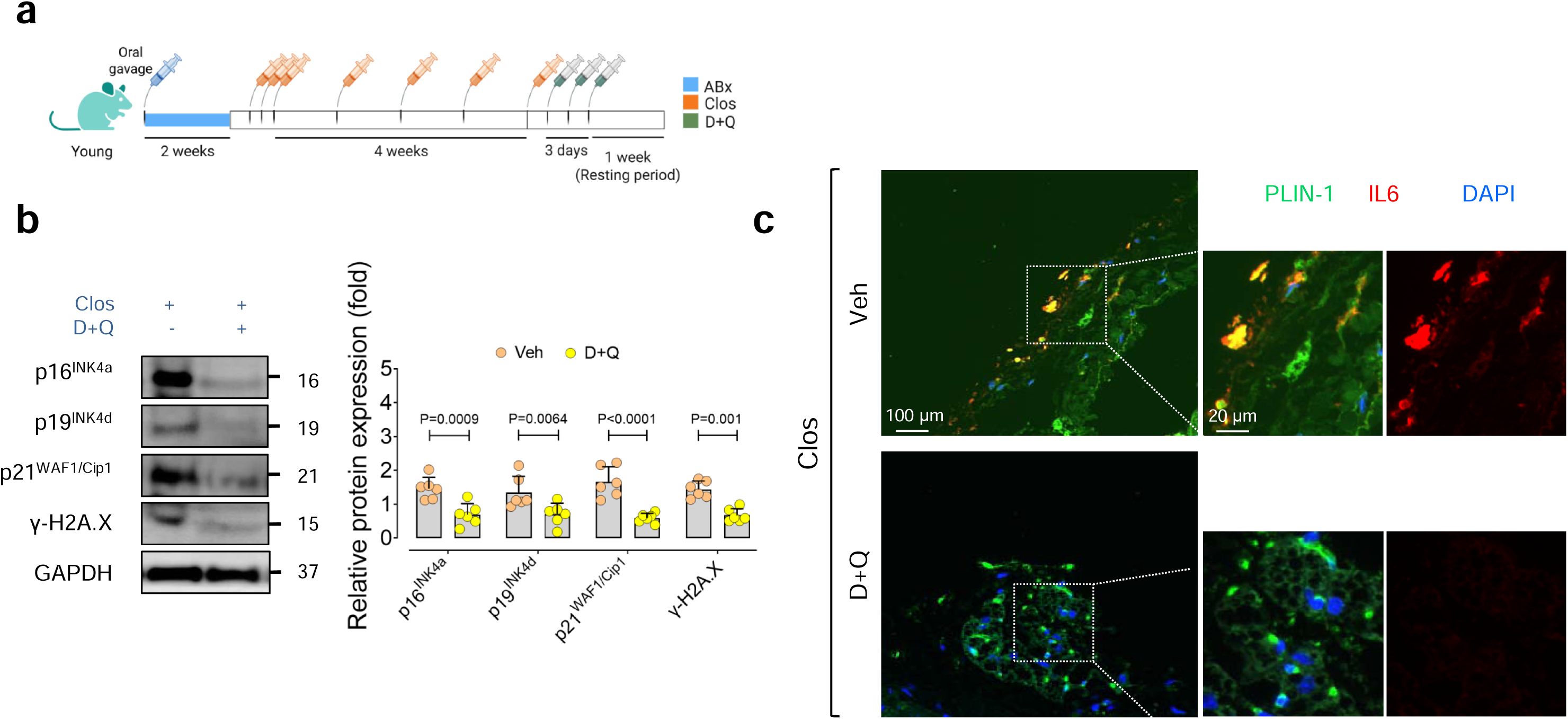
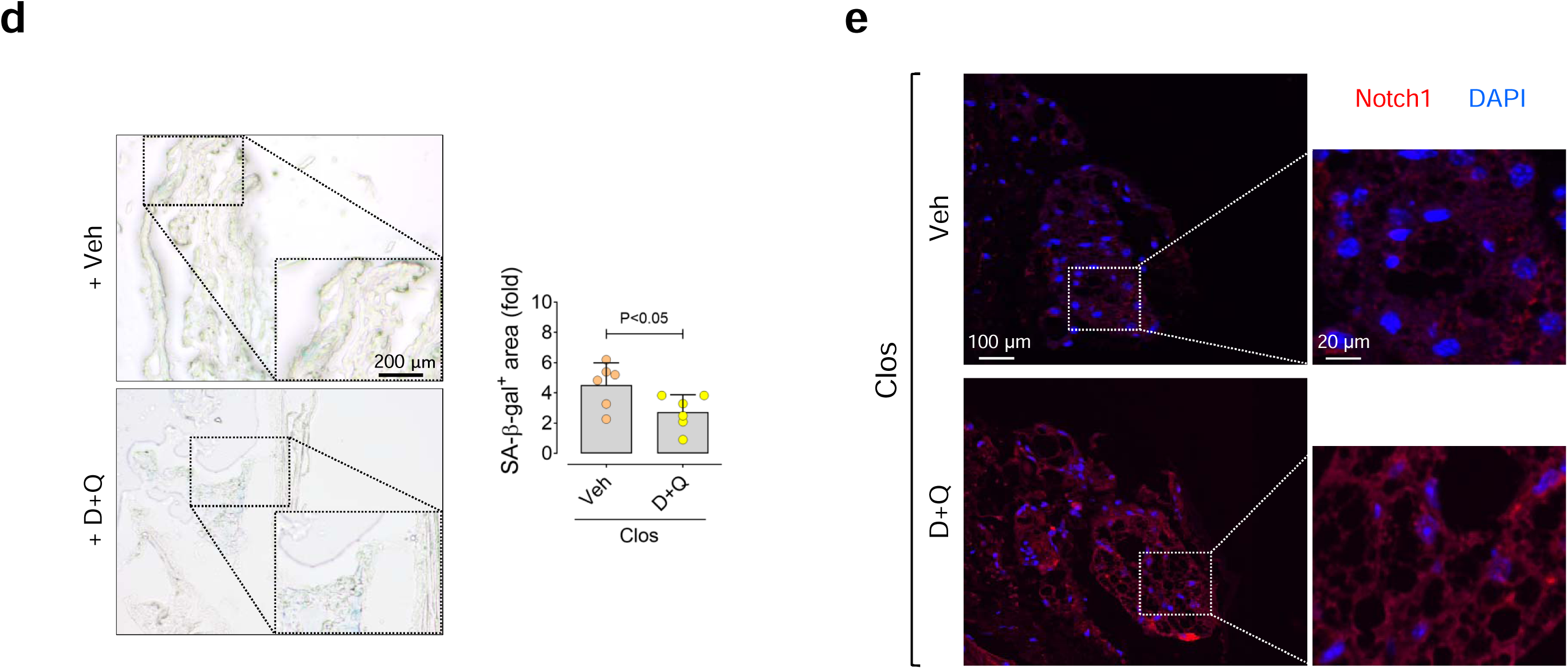
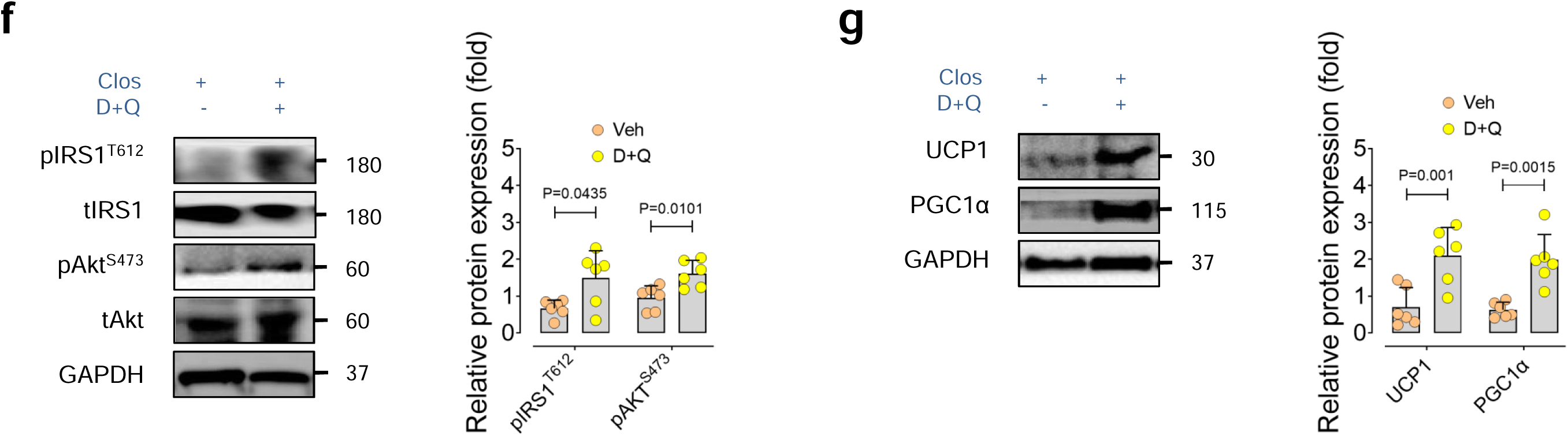
Senolytics rescue PAA-induced PVAT senescence. **a,** *Clos*-colonized young mice received a senolytic cocktail containing Dasatinib + Quercetin (5 + 50 mg/kg/d) for 3 days, followed by a 7-day resting period (*n*□=□6). **b,** Representative immunoblots for CDK inhibitors p16^INK4A^, p19^INK4D^, and p21^WAF1/Cip1^ and DNA damage marker γ-H2A.X in tPVAT from *Clos*-colonized mice treated with D+Q or vehicle (*n* = 6). **c,d,** Representative confocal immunofluorescence images of IL-6 (**c**) and bright-field SA-β-gal staining images (**d**) in tPVAT from these mice (*n* = 6). **e,** Representative confocal images of NOTCH1 in tPVAT from these mice (*n* = 6). **f,g,** Immunoblotting for the insulin signaling pathway (**f**) and thermogenic markers UCP1 and PGC1α (**g**) in tPVAT from *Clos*-colonized mice treated with D+Q or vehicle (*n* = 6). Data were determined in 8-10 micrographs and represent triplicated biologically independent experiments. Scale bar, 50, 100, and 200 μm (**c,d,e**). Error bars represent SD (**b,d,f,g**). *P* values were calculated using a two-tailed unpaired Student’s *t*-test (**b,d,f,g**). Images created with https://BioRender.com (**a**).

At the systemic level, D+Q-treated *Clos*-colonized mice exhibited improved metabolic health, reflected by reduced body weight (mean 31.87 *vs.* 24.67□g) and lower serum insulin levels (Extended Data Fig. 8a,b). As a proof of concept, we employed p16-3MR mice, which display vascular and PVAT senescence phenotypes closely resembling those observed in *Clos*-colonized mice (Extended Data Fig. 9a–d). Our data revealed that pharmacological clearance of senescent cells using GCV significantly downregulated pro-atherogenic and tissue-remodeling genes (*Il1b*, *Il6*, *Tnfa*, *Vcam1*, *Mmp2* and *Mmp9*) in aortic tissues (Extended Data Fig. 9e). Concurrently, GCV treatment enhanced the expression of thermogenic and anti-inflammatory genes (*Ucp1* and *AdipoQ*) while suppressing pro-inflammatory mediators (*Il1b*, *Il6*, *Ccl2*, *Tnfa* and *Leptin*) in tPVAT (Extended Data Fig. 9f).

Together, these findings strongly confirm first, the direct contribution of *Clostridium* sp. ASF356 to PVAT senescence and dysfunction and, second, the potential of the reversal of PVAT dysfunction and insulin resistance by senolytic therapy *in vivo*.

### PAA actively contributes to PVAT dysfunction and atherosclerosis

Vascular ECs and PVAT engage in a dynamic paracrine crosstalk that plays a critical role in vascular homeostasis and atheroprevention. Cellular senescence and dysfunction in either compartment disrupt this homeostatic interaction, fostering a pro-inflammatory vascular milieu and reinforcing atherogenesis^20–22^. Given our previous findings and the present mechanistic evidence demonstrating that PAA triggers senescence and dysfunction in both ECs and PVAT, we hypothesized that elevated PAA levels causally contribute to atherosclerosis development. To address this, we first quantified plasma PAA concentrations by targeted LC-MS/MS in an ASCVD cohort (*Aging Heart Zurich Cohort*), comprising 110 aged patients (> 80 years old) with ASCVD and 77 non-ASCVD control individuals (Fig. 6a; Supplementary Table 1). As shown in Fig. 6b, PAA levels were significantly higher in ASCVD patients than in control counterparts (median [IQR]: 144.6 [93.5–221.3] µM *vs.* 111.0 [54.1–170.0] µM). Stratification by tertiles further revealed that individuals in the highest PAA tertile (T3) exhibited a markedly higher ASCVD prevalence compared with those in the lowest tertile (T1) (Fig. 6c; Supplementary Table 2). Importantly, multivariable logistic regression model adjusting for traditional risk factors showed that elevated PAA levels remain independently associated with ASCVD, supporting PAA as an indicator of increased atherosclerotic risk. Consistent with these clinical observations, atherosclerosis-prone *Ldlr^-/-^* mice fed a Western diet (WD; high-cholesterol diet) for 12 weeks exhibited significantly higher PAA levels compared to wild-type mice maintained on either chow or Western diet (Fig. 6d,e). Our H&E histological analyses also confirmed pronounced acceleration of aortic root plaque progression in WD-fed *Ldlr^-/-^* mice (Fig. 6f). Notably, we identified PAA concentrations strongly correlated with both aortic root lesion area (*R* = 0.3832, *p* = 0.0036) and luminal occlusion (*R* = 0.3921, *p* = 0.0031) in these mice (Fig. 6g), reinforcing a robust association between PAA and atherosclerosis burden.

**Fig. 6.**
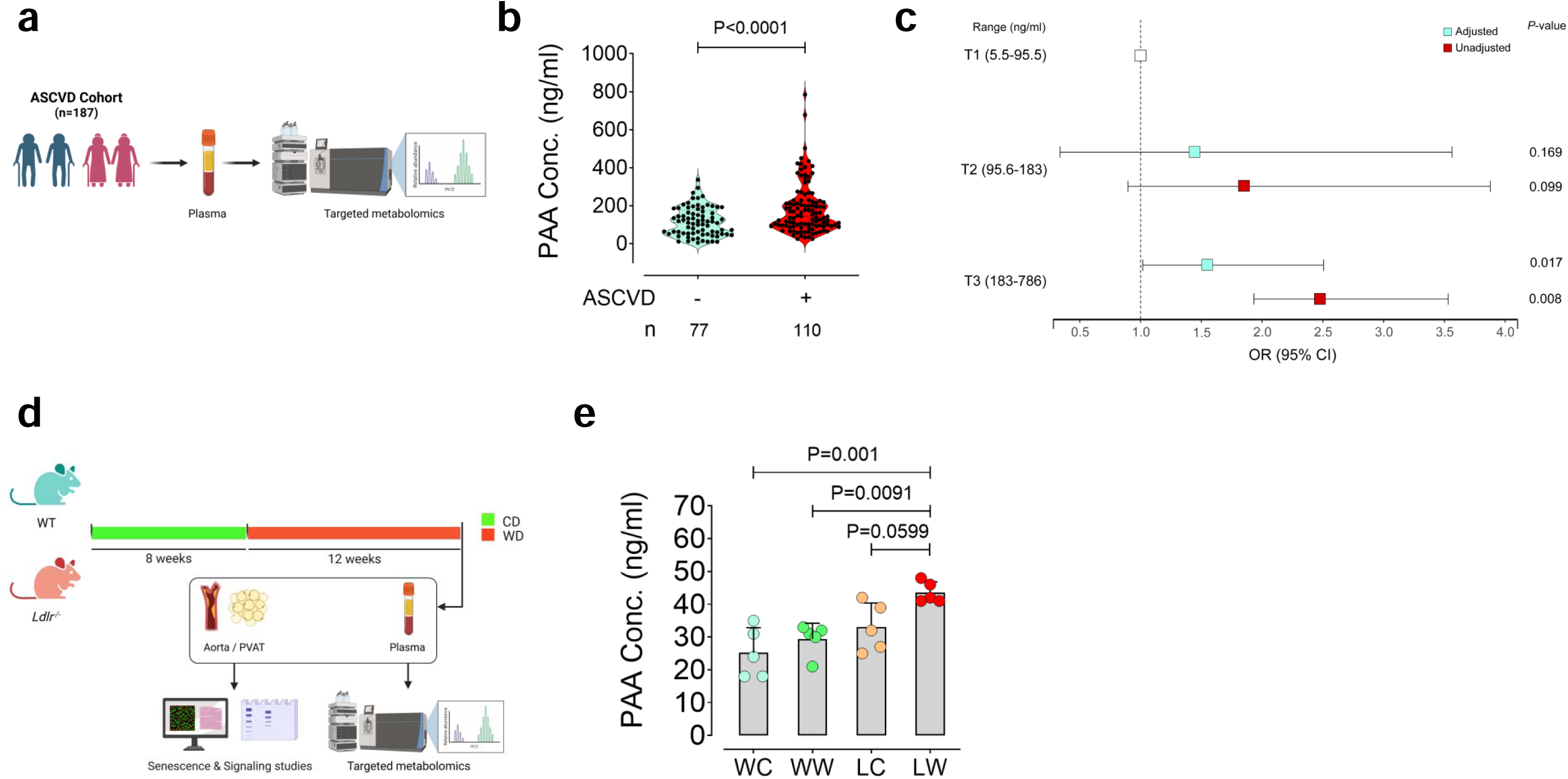
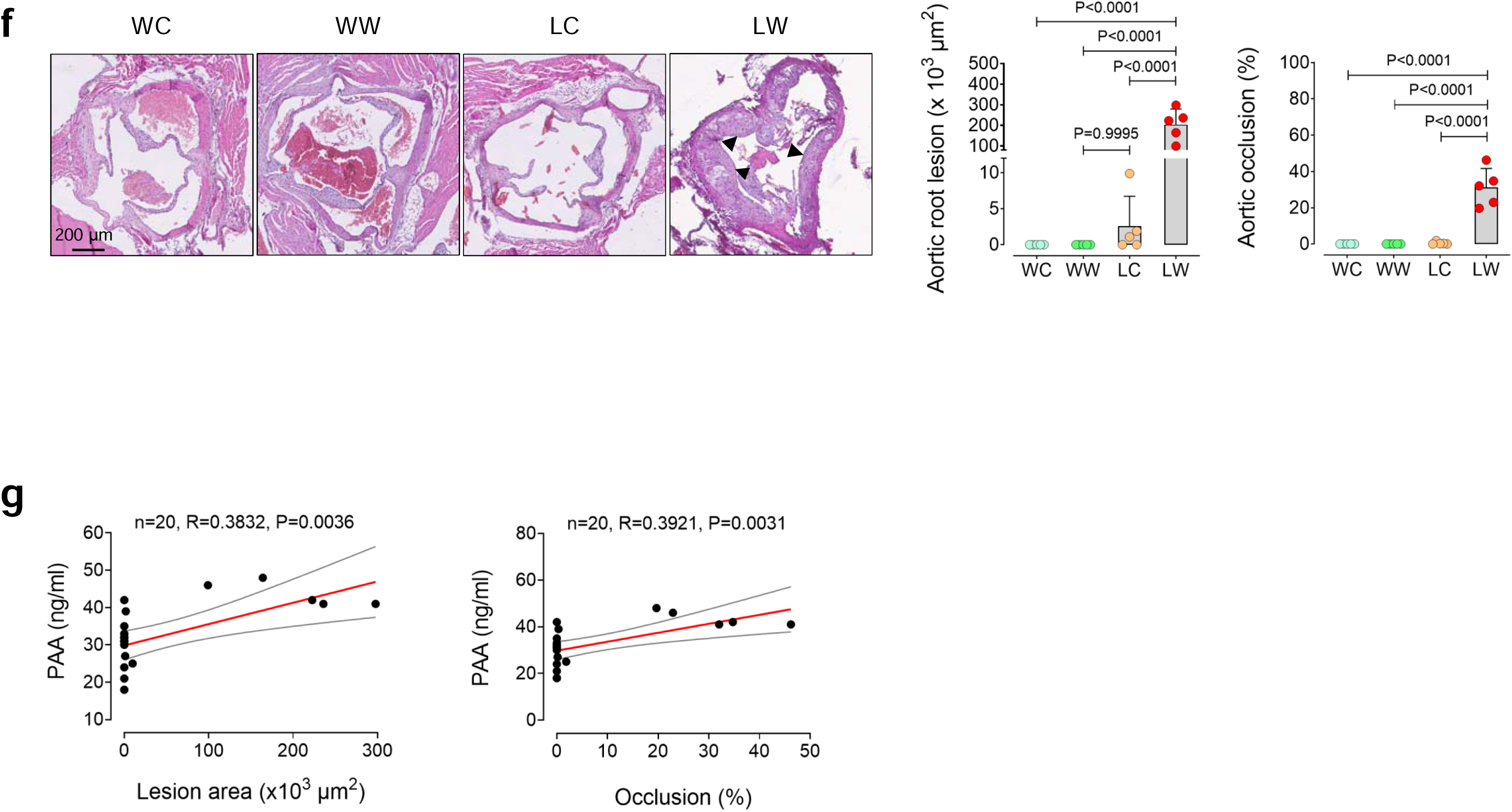
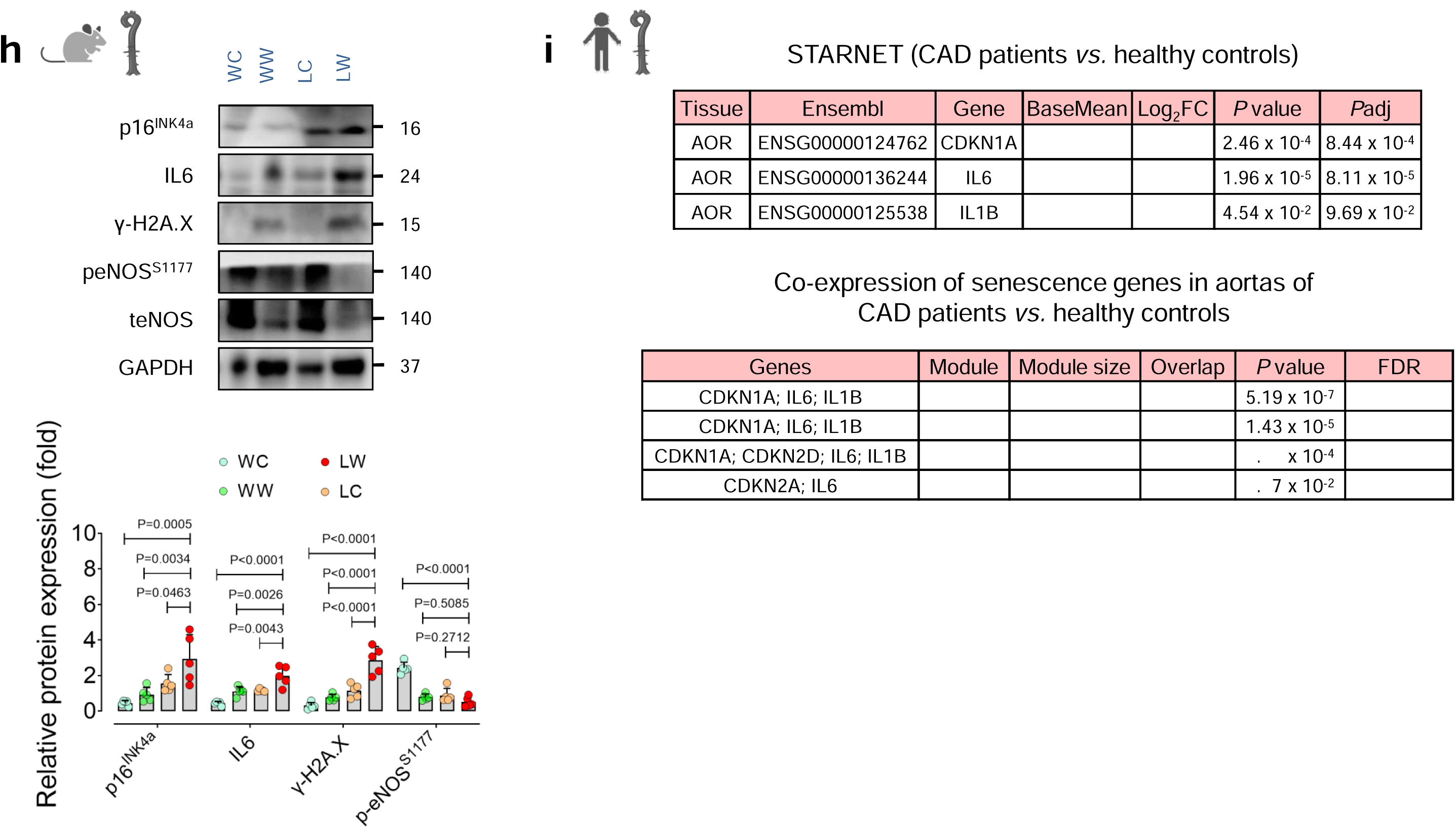
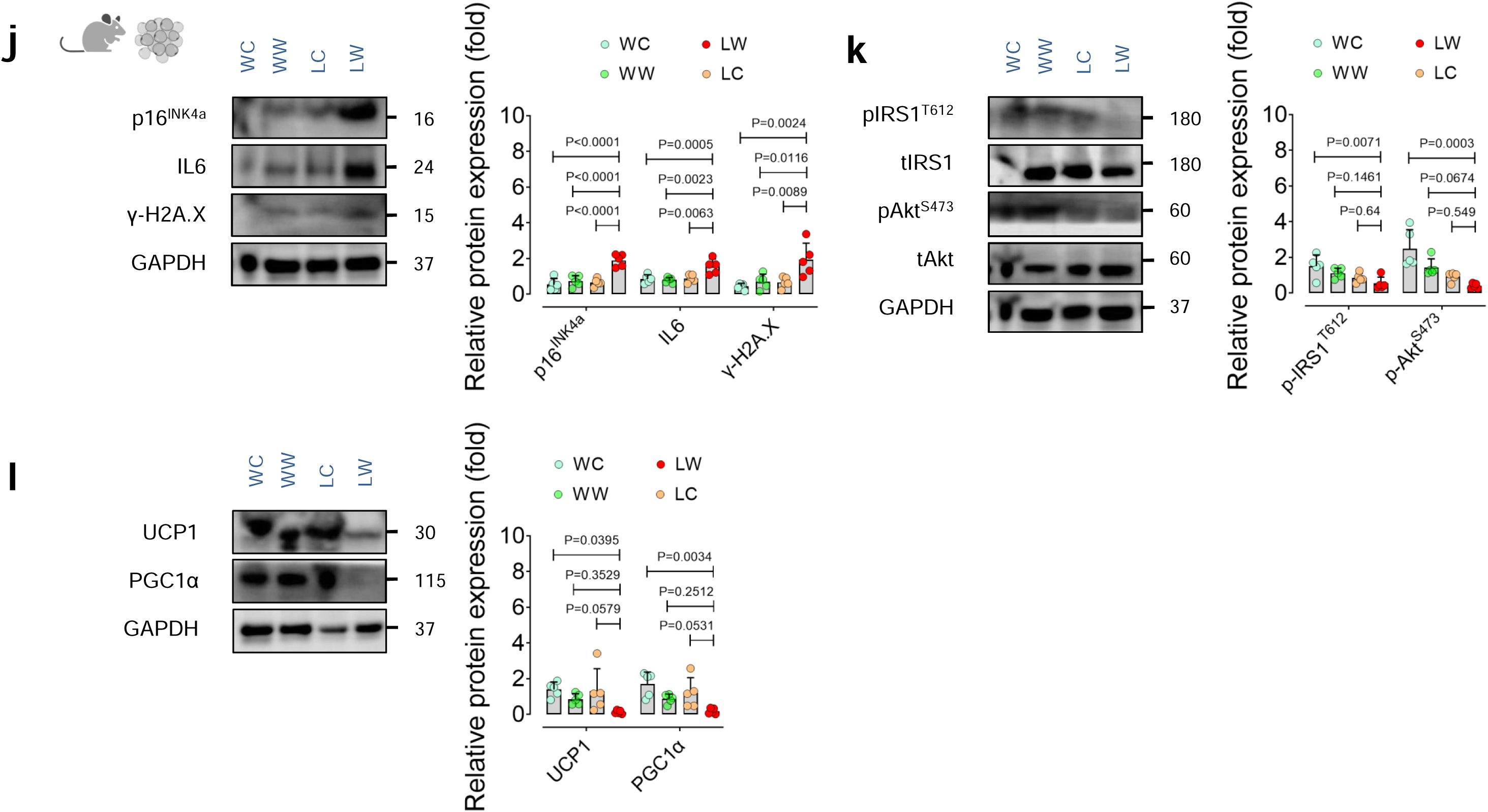
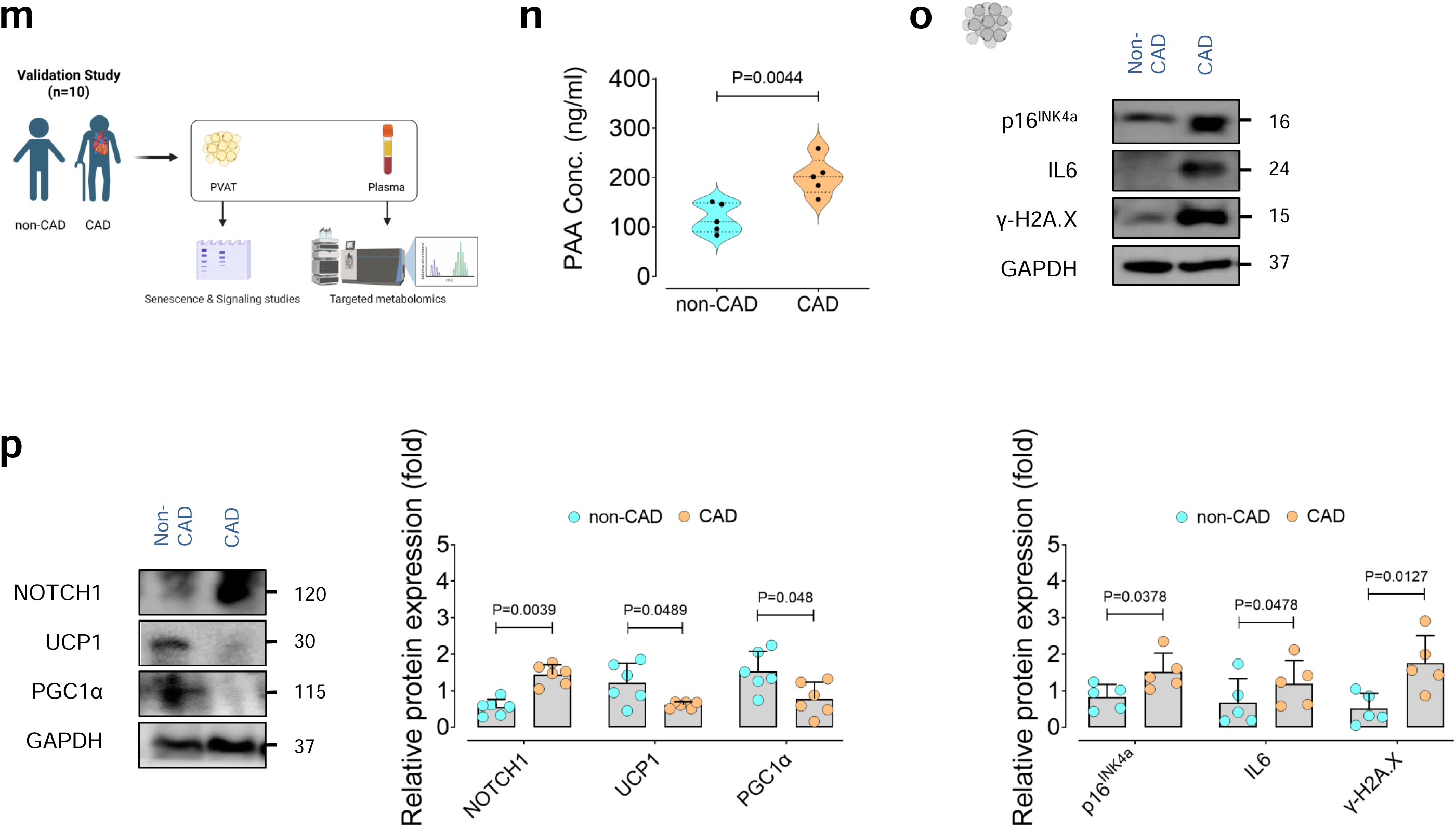
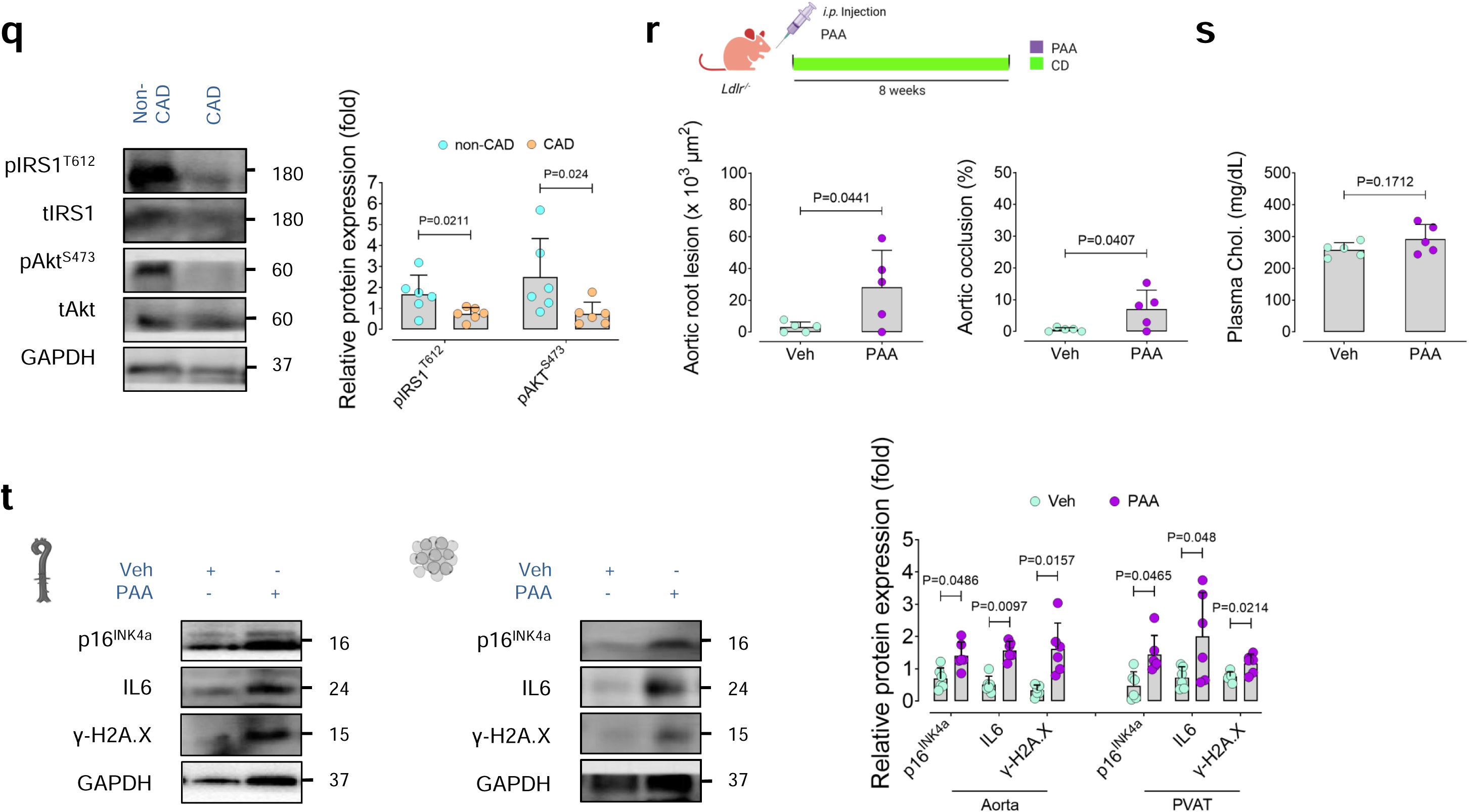
PAA contributes to PVAT dysfunction and is linked to atherosclerosis. **a,b,** Plasma samples from aged ASCVD patients (>80 years old) enrolled in the ASCVD cohort (*n* = 110; male and female) and healthy controls (*n* = 77; male and female) were subjected to targeted metabolomics for PAA quantification. **c,** Adjusted regression models for the association of PAA with atherosclerosis in the ASCVD cohort (*n*□=□187). Effect estimates were controlled for age, sex, smoking, alcohol, family history of CVD, LDL-C, triglycerides, HDL-C, Hb1Ac, CRP, troponin T, and NT-proBNP. Error bars show 95% confidence intervals. OR, odds ratio. **d,** *Ldlr^−/−^* and WT mice were fed chow (for 8 weeks) or Western diet (for 12 weeks). WC: WT + chow diet; WW: WT + Western diet; LC: *Ldlr^−/−^* + chow diet; LW: *Ldlr^−/−^* + Western diet. **e,** Plasma PAA levels in these mice quantified by LC-MS/MS targeted metabolomics (*n*□=□5). **f,** Representative images (left) and quantification (right) of H&E staining of aortic root lesions (*n*□=□5). Arrowheads indicate plaque areas. **g,** Correlation of plasma PAA levels with aortic root lesion area and luminal occlusion (%). **h,** Representative immunoblots and quantification of intensities for the CDK inhibitor p16^INK4A^, the SASP component IL-6, DNA damage marker γ-H2A.X, and the endothelial function marker phosphorylated eNOS^S1177^ in aortas from *Ldlr^−/−^* and WT mice fed a chow or Western diet for 12 weeks (*n* = 5). **i,** Expression of *CDKN1A*, *IL1B*, and *IL6* genes (upper) and co-expression of the senescence-associated genes in atherosclerotic aortic wall (AOR) of patients with CAD (*n*□=□600) and healthy individuals (*n*□=□250) in the STARNET database. **j,k,l,** Representative immunoblots and quantification of intensities for senescence hallmarks (**j**), insulin signaling pathway (**k**), and thermogenic markers (**l**) in tPVAT from *Ldlr^−/−^* and WT mice fed a chow or Western diet for 12 weeks (*n* = 5). **m,** Plasma and aortic PVAT samples were collected from the validation study, including aged CAD patients undergoing CABG surgery (*n*□=□5) and non-CAD controls (*n*□=□5), for LC-MS/MS PAA quantification and senescence studies. **n,** Plasma PAA concentrations in aged CAD patients (*n*□=□5) and non-CAD controls (*n*□=□5). **o,p,q,** Representative immunoblots and quantification of intensities for senescence hallmarks (**o**), NOTCH1 and thermogenic markers (**p**), and insulin signaling pathway (**q**) in aortic PVAT from the individuals in the validation study (*n* = 5). **r,** PAA was administered (PAA) or not (Ctrl) to chow-fed *Ldlr^−/−^* mice for 8 weeks. **s,** Quantification of H&E-stained aortic root lesion area (left) and aortic occlusion (right). Total cholesterol concentrations in plasma (*n* = 5). **t,** Representative immunoblots and quantification of intensities for senescence hallmarks p16^INK4A^, IL-6, and γ-H2A.X in aortas (left) and tPVAT (right). Data were determined in 8-10 micrographs and represent triplicated biologically independent experiments. Scale bar, 200 μm (**f**). Error bars represent SD (**h,j-l,o-t**). *P* values were calculated using two-tailed Mann–Whitney *U*-test (**b**), one-way ANOVA followed by Tukey’s post *hoc* test (**e,f,h,j-l**), Welch’s t-test (**i**), and a two-tailed unpaired Student’s *t*-test (**n-t**). Correlation coefficient and *P* values were calculated by Spearman’s rank-order correlation test (**g**). Data are shown as median with min–max; each violin represents interquartile range (IQR); center lines indicate the median; upper and lower lines are bounded by 25th and 75th percentiles (**b**,**n**). Images created with https://BioRender.com (**a,d,m,r**).

Given that PAA robustly accelerates vascular senescence, we hypothesized that its pro-senescent activity constitutes a mechanistic driver of atherosclerosis development. We therefore characterized senescence phenotype in aortic ECs, which we earlier in this study demonstrated to propagate senescence and dysfunction to adjacent PVAT. In WD-fed *Ldlr^-/-^*mice, we observed a marked upregulation of CDK inhibitor p16^INK4A^, the SASP component IL6, and DDR marker γ-H2A.X in aortic ECs. These changes coincided with downregulated eNOS^S1177^ phosphorylation, a key determinant of endothelial function^2,23,24^ (Fig. 6h). To extend these findings to human disease, we further interrogated senescence-associated gene expression in aortic tissues from patients with coronary artery disease (CAD; *n*□=□600) and healthy individuals (*n*□=□250) from the Stockholm-Tartu Atherosclerosis Reverse Network Engineering Task (STARNET) database^25^. Consistent with our experimental data, the expression of individual senescence-related genes, including *CDKN1A*, *IL6*, and *IL1B*, as well as their co-expression network, was significantly higher in atherosclerotic aortic tissues from patients with CAD compared with healthy controls (Fig. 6i). These findings support a conserved senescence-driven transcriptional program in atherosclerotic vasculature and reinforce a mechanistic link between PAA-induced endothelial senescence and human atherosclerosis.

In line with our earlier identification of endothelial senescence as a driver of the aortic inflammatory milieu, we found that tPVAT surrounding atherosclerotic aortas from WD-fed *Ldlr^-/-^* mice exhibit marked elevations in key senescence hallmarks p16^INK4a^, IL-6, and γ-H2A.X (Fig. 6j), together with impaired insulin signaling (Fig. 6k). In parallel, thermogenic programming was suppressed in tPVAT, as evidenced by downregulation of UCP1 and PGC1α (Fig. 6l). Consistently, these mice displayed a reduced insulin sensitivity phenotype compared with WT controls, characterized by significantly higher serum insulin levels, increased body weight, and elevated plasma cholesterol and triglycerides (Extended Data Fig. 10a–c), underscoring the contribution of PAA to insulin resistance and metabolic dysfunction.

As a proof of concept in humans, we next examined a validation study on aortic PVAT obtained from aged CAD patients undergoing CABG surgery, who exhibited elevated plasma PAA concentrations compared with non-CAD controls (Fig. 6m,n; Supplementary Table 3). Our findings showed senescence phenotype, represented as upregulated p16^INK4A^, γ-H2A.X, and IL-6 in tPVAT specimens from CAD patients (Fig. 6o), mirroring senescence-associated PVAT dysfunction in atherosclerosis-prone mice. In line with these findings, NOTCH1 expression was robustly increased, concomitant with suppression of adipocyte thermogenic markers UCP1 and PGC1α (Fig. 6p), where insulin signaling was impaired (Fig. 6q) in CAD patients.

To further determine whether PAA causally contributes to early stages of atherosclerosis, we administered chow-fed *Ldlr^−/−^* mice to PAA for 8 weeks. PAA supplementation markedly accelerated atherosclerosis development in the aortic root, as evidenced by significant increases in atherosclerotic lesion area (∼9.1-fold) and luminal occlusion (∼12-fold) (Fig. 6r), without affecting circulating total cholesterol levels (Fig. 6s). Of note, the expansion of atherosclerotic plaque burden was accompanied by robust induction of senescence hallmarks p16^INK4A^, IL-6, and γ-H2A.X in both aortic ECs and tPVAT, thereby establishing a pro-atherogenic vascular microenvironment (Fig. 6t; Extended Data Fig. 11).

Collectively, these data identify PAA as a novel contributor to atherosclerosis aetiology that acts independently of circulating cholesterol concentrations. Instead, the pro-atherogenic effects of PAA are mediated through its capacity to induce cellular senescence within the (peri)vascular niche, thereby coupling microbial metabolism to vascular aging and disease progression.

## Discussion

The complex aetiology of ASCVD poses challenges to the detection of early stages of the disease and novel therapeutic strategies^3,6,26,27^. Previous studies have focused on associations of gut microbiota-dependent metabolites with advanced stages of ASCVD or mortality^5,28–31^. Our study identifies the gut microbiota–generated metabolite PAA as a modifiable risk factor for ASCVD in ageing. It establishes a direct link between age-dependent increases in circulating PAA and PVAT senescence, revealing PVAT as an unprecedented downstream target of microbiota-driven ageing signals in the host. We previously demonstrated that members of the *Clostridium* genus—particularly *Clostridium* sp. ASF356—are enriched in the aged microbiome and generate PAA from phenylalanine, thereby inducing aortic endothelial senescence and vascular dysfunction^2^. Building on this work, we now show that age-associated elevations in circulating PAA initiate a paracrine senescence-promoting cascade, in which senescent endothelial cells propagate dysfunction to adjacent PVAT. Together, these findings expand the gut–vascular axis to encompass endothelial–adipocyte crosstalk as a central determinant of age-accelerated atherosclerosis.

Using human cohort data and experimental mouse models, this work reveals that PAA-induced endothelial senescence triggers SASP enriched in IL-6, which in turn activates NOTCH1 signaling in PVAT adipocytes. It was previously demonstrated that the stimulation of NOTCH1 and its downstream target Hes1 represses beige and thermogenic genes including *Ppargc1*α and *Prdm16*, alongside *Ucp1* downregulation^16,32^. In obese humans and mice fed a high-fat diet, NOTCH1 activation also exacerbates insulin resistance accompanied by marked alterations in the pro-inflammatory secretory profile and downregulated mitochondrial respiration and ATP biosynthesis in PVAT^16,19,32–34^. Our findings demonstrate this endothelial-adipocyte axis disrupts insulin signaling and thermogenic programming, culminating in a dysfunctional PVAT phenotype that is increasingly recognized as a pro-atherogenic factor^12,21,35,36^. These results align with prior work showing that senescence-messaging secretomes from senescent endothelial cells induce a senescence-like state and impair insulin signaling in adipocytes, thereby promoting metabolic dysfunction in EC-specific progeroid mice^9^. Conversely, selective elimination of p16^INK4a+^ senescent endothelial cells attenuates the SASP in adipose tissues and restores metabolic homeostasis in obese *Tie2-Cre*, *p16^Ink4a^-LOX-ATTAC* mice^10^. Importantly, our data establish that microbial metabolites can exert indirect yet durable effects on vascular homeostasis by reshaping the local cellular niche, rather than acting solely through direct endothelial- or immune-mediated mechanisms^37^. In this context, PVAT emerges as an active amplifier of microbial aging signals, integrating metabolic and inflammatory cues to influence atherosclerosis progression^12,21^. The translational relevance of this pathway is underscored by our analyses of PVAT obtained from aged CAD patients undergoing CABG surgery. Elevated plasma PAA levels were associated with a senescent, metabolically suppressed PVAT phenotype, closely mirroring our experimental observations. Notably, selective clearance of senescent cells using the senolytic D+Q cocktail restored PVAT function, improved insulin signaling, and attenuated metabolic dysfunction in mice. In line with this, our data demonstrated that elimination of senescent cells alleviates pro-atherogenic and pro-inflammatory transcriptional changes in p16-3MR mice. These findings provide direct causal evidence that PAA-induced senescence is not merely correlative but functionally contributes to vascular pathology, and that targeting senescent cell burden represents a viable strategy to mitigate microbiota-driven vascular aging.

These findings were independently validated in an ASCVD cohort, further positioning PAA as a companion diagnostic biomarker of atherosclerosis and highlighting its potential as a therapeutic target for personalized medicine. Moreover, we demonstrate that PAA drives atherosclerosis independently of blood cholesterol concentrations. Pro-atherogenic effects of PAA can be attributed to its senescence-promoting effects across multiple cell types including endothelial cells and adipocytes. We therefore observed that administration of PAA induced senescence—characterized by cell-cycle arrest, SASP, and DNA damage response—in aortic endothelial cells and PVAT, thereby contributing to atherosclerosis progression and further plaque instability. Although PAA administration did not significantly alter cholesterol levels or key renal and hepatic function parameters, we cannot rule out additional systemic effects that may contribute to the disease pathology and warrant future studies.

The discovery of a direct link between gut microbiota-mediated Phe metabolism and ASCVD risk takes another step forward toward uncovering the full picture of the gut-vascular axis in ageing that potentially pinpoints novel therapeutic targets for broader healthy ageing implications. Our findings uncover a previously unrecognized mechanism by which PAA triggers senescence phenotype within the aortic endothelial-adipocyte microenvironment, thereby linking with atherosclerosis progression. The role of *Clostridium* species in this senescence-promoting pathway highlights new opportunities for microbiome-based senotherapies, such as bacterial enzyme inhibition or genetic engineering, to prevent ASCVD in aging populations. Given that ASCVD event rates remain high despite optimal management of traditional risk factors, novel microbiome-targeted interventions may offer a complementary and potentially more effective approach to risk reduction^6,38–40^. Beyond its pathogenic role, our data suggest that circulating PAA may serve as a biomarker of vascular aging, with potential utility for predicting premature ageing process across multiple body organs. This work also raises important considerations regarding the safety of high dietary intake of Phe-rich foods, which under certain microbiome configurations may foster the expansion of PAA-producing bacteria and enhance the generation of downstream metabolites such as PAGln—a metabolite previously associated with major adverse cardiovascular events in patients with cardiometabolic diseases^7,15,41–43^. Such metabolic reprogramming may thereby accelerate vascular aging and atherosclerosis. Taken together, our study revealed that vascular senescence triggered by gut-derived PAA is a bona fide non-traditional risk for ASCVD in aging populations, and thus provides scientific proof for the famous saying: “a man is as old as his arteries”.

## Methods

### Human cohorts

#### – TwinsUK cohort

To investigate associations between circulating PAA concentrations and age, we used metabolomic data from the TwinsUK human aging cohort^44^. This cohort comprises >14,000 predominantly same-sex adult twins aged 18–95 years, of whom a subset underwent metabolomic profiling. All participants received no compensation and provided written informed consent. The study was approved by the NRES Committee London-Westminster (REC Reference No.: EC04/015).

Metagenomic data from the same cohort were used to examine associations between PAA and either *ppfor* (pyruvate:ferredoxin oxidoreductase; KEGG Orthology K00179) abundance or specific species. Paired-end fecal metagenomes (>10M reads per sample) were processed with YAMP v0.9.5. Taxonomic profiling and relative abundance of all bacterial species were quantified using MetaPhlAn v4.0^45^, using an updated species-specific marker database built from 99,237 reference genomes representing 16,797 species (GenBank, January 2019). Metagenomes were mapped internally in MetaPhlAn v.4.0 against the marker gene database with Bowtie 2 v.2.3.4.3 in “very-sensitive” mode, and alignments with MAPQ < 5 were excluded to remove low-confidence matches.

Functional profiling was conducted using HUMAnN3 pipeline for associations between relative *ppfor* abundance (copies per million) and PAA. The abundance values were kept untransformed, and outliers >3 standard deviations from the group mean were excluded prior to association analyses.

#### – ASCVD cohort

To examine the associations between plasma PAA levels and atherosclerotic CVD, we collected plasma samples from aged patients (>80 years old) with ASCVD (*n* = 110) enrolled in *Aging Heart Zurich Cohort—* an ongoing observational prospective human cohort study conducted at University Hospital Zurich^46^ (Supplementary Tables 1 and 2). All patients were diagnosed for ASCVD, including atherosclerosis, coronary artery disease, and arrhythmias, at baseline in 2020 and are followed up every 1 year. Importantly, individuals with known intestinal disorders or who had taken antibiotics or consumed alcohol within the three months prior to sample collection were excluded. Atherosclerosis was assessed by imaging studies cardiac CT scan and coronary angiography. Fasting blood test was performed for biochemical parameters including blood glucose, Hemoglobin A1c (HbA1c), total cholesterol, LDL-cholesterol, HDL-cholesterol, triglyceride, CRP, BUN, creatinine, alanine aminotransferase (ALT), aspartate aminotransferase (AST), and gamma-glutamyl transferase (GGT), troponin T, and NT-proBNP. All participants received no compensation and provided written informed consent. The study was approved by the Kantonale Ethikkomission Zurich (BASEC-Nr: 2020-02597). Plasma samples from a separate cohort of 77 healthy controls were obtained from the blood bank of the Swiss Red Cross.

#### – Validation cohort

To validate the associations between plasma PAA levels and age in healthy humans, we obtained plasma samples from a cohort of 98 healthy subjects from the blood bank of the Swiss Red Cross and processed accordingly.

### Human aorta and PVAT explants

CAD Patients (*n* = 5) were undergone to coronary artery bypass graft (CABG) surgery at Department of Cardiac Surgery, University Hospital Zurich, Switzerland (Supplementary Table 3). All procedures were approved by the local ethics committee and performed in accordance with the Declaration of Helsinki (Kantonale Ethikkommission Zürich, BASEC number: 2023-02040). Thoracic PVAT tissues (from paracardial region) were collected and snap frozen in liquid nitrogen for senescence and signaling studies. Tissue samples from non-CAD transplant donors (*n* = 5) served as controls. Plasma samples were also stored at - 80□C for further LC-MS/MS quantification of PAA.

### Mouse models

Aging experiments were performed using wild-type C57BL/6J female and male mice purchased from the Jackson Laboratory at the age of 8 weeks and aged in the animal facility of University of Zurich, Schlieren. Accordingly, mice were fed a standard chow diet (containing 19% Protein, 61% Carbohydrate and 7% Fat, #D11112201, Research Diets, New Brunswick, NJ, USA) until 3 (as Young) or >24 (as Old) months of age. Our study complies with all ethical regulations approved by the Institutional Animal Care and Use Committee at University of Zurich and the Cantonal Ethics Commission Zurich (ZH023/2023 and ZH241/19).

Atherosclerosis-prone *Ldlr*-KO C57BL/6J mice were fed a specific Western diet (21% fat, 0.21% cholesterol, SSNIFF TD. 88137, E15775-34; Supplementary Table 4) or standard chow diet (as controls) for 12 weeks, as indicated in each experiment. PAA (50 mg/kg per day, *i.p.*, Sigma-Aldrich, P16621) was administered for 8 weeks. Mice were kept at the animal facility of the University of Milan, Italy. All animal procedures were approved by the pertinent Ethical Committees (Autorizzazione Ministeriale 384/2019-PR, 92/2020, 240/2021 and 565/2022).

The p16-3MR mice were kept under specific pathogen-free conditions at the animal facility of the Institute for Research in Biomedicine (IRB), Università della Svizzera Italiana, Bellinzona, Switzerland. Mice were administered with vehicle (PBS) or GCV (25 mg/kg) via daily *i.p.* injection for 5 consecutive days. The sequences of primers used for PCR genotyping are provided in Supplementary Table 5. Positive mice demonstrated a band at 202□bp. Experiments are approved by the local ethics committee (the local ethics committee, Canton Ticino), authorization numbers 30275 and 34293.

Animal care and all experimental protocols were in accordance to the principles of the ‘3Rs’—replacement, reduction and refinement—according the Directive 2010/63/EU of the European Parliament and of the Council of 22 September 2010 on the protection of animals used for scientific purposes. All mice were housed in controlled environment in plexiglass cages under a strict 12 h:12 h light/dark cycles at an ambient temperature (23 ± 1°C) and humidity of 55 ± 10%, and had free access to drinking water and food *ad libitum*.

### Blood and fecal collection, and tissue harvesting

Feces were collected from distal colon, quickly snap-frozen, and stored at -80°C for DNA extraction and shotgun metagenomic sequencing.

Blood samples were obtained by cardiac puncture. The plasma and serum were collected by centrifugation of aliquots of blood samples at 125 x g (8 min) and 1500 x g (10□min) at room temperature and frozen at −80□°C^2,47^. After blood collection, heart was perfused for aorta and PVAT harvesting.

### Targeted LC–MS/MS analysis of plasma PAA levels

PAA levels in human and mouse plasma (100 μL) were quantified by liquid chromatography coupled to a high resolution mass spectrometer (LC-HRMS), as previously described^2^. Plasma was extracted with frozen ethanol/methanol (-20°C) containing internal standard (D5-PAA, 0.25 µg/mL) and centrifuged (13,000 rpm, 4°C, 20 min). Supernatants were reconstituted into 100µl of water/methanol (90/10) for next centrifugation. Supernatants were separated in a Waters HSST3 column (150 × 2.1 mm, 1.8 μm) and analyzed on a Thermo Ultimate 3000 LC system coupled to a Thermo Exploris 120 orbitrap mass spectrometer. Peaks were identified using standards and processed with Tracefinder 3.2 (Thermo Scientific). All MS conditions are detailed in Table 1.

**Table 1.** Plasma PAA mass spectrometry conditions.

|  | Polarity | Formula | Adduct | Precursor ion (m/z) | Product ion (m/z) | HCD (%) |
| --- | --- | --- | --- | --- | --- | --- |
| PAA | Negative | $\text{C}_8\text{H}_8\text{O}_2$ | -H | 135.0452 | 91.0553 | 30 |
| PAA-d5 | Negative | $\text{C}_8\text{H}_3\text{O}_2\text{D}_5$ | -H | 140.0765 | 96.0869 | 30 |

### Fecal DNA extraction

Fecal genomic DNA was extracted using the FastDNA™ SPIN kit (MP Biomedicals, 116540600-CF)^2,47^. Samples were homogenized in MT buffer with FastPrep, centrifuged, and the supernatant was mixed with PPS buffer. DNA was purified using a column, dissolved in sterile Milli-Q water, and assessed by a NanoDrop 2000 (Thermo Scientific).

### Shotgun metagenomics

Metagenome libraries were generated using a high-quality genomic DNA and libraries were sequenced on an Illumina NovaSeq 6000 PE150 platform (Novogene, UK) with a 250-bp paired-end model.

#### Taxonomic profiling

Taxonomic profiling was performed as previously described^2^. Briefly, we used kraken2/Bracken2 (parameters: *--confidence 0.51*)^48^ against the mouse microbiota genome catalog (CMMG) (https://ezmeta.unige.ch/CMMG/Kraken2db/cmmg)^49^. Kraken2 profiles were then subjected to Bracken2 for taxonomic refinement using the corresponding database and read length. Bracken2 corrected CMMG read counts were then normalized for reference genome length and loaded together with available reference genomes taxonomy (https://ezmeta.unige.ch/CMMG/Supplementary_Tables_and_Figures/Tables/Table_S4_curated_taxonomy.tsv) into a phyloseq object for data handling. Only the taxa identified in at least 2 samples with coverage of at least 100 reads in total were considered for further analyses.

#### Functional profiling

Genomes were screened for presence of K00169 (*vor*) and or K00179 (*ppfor*), the genes encoding the enzymes participated in the phenylalanine metabolism and production of PAA, based on their available Kyoto Encyclopedia of Genes and Genomes (KEGG) functional annotations (https://ezmeta.unige.ch/CMMG/functional_annotations/mouse/annotations.tsv). The differential abundance of K00169 or K00179 in the identified genome was then added as additional information in the phyloseq object to identify VOR*-* and/or PPFOR-positive taxa in the microbiome of old and young mice. The association between the abundance of KO and plasma levels of PAA were also characterized by a Spearman correlation analysis.

#### Community analysis

Microbiota profiles were characterized using alpha-diversity and beta-diversity^2,47^. Hill-d0, 1, and 2 indices represent the richness of detected taxa in aged or young microbiome. Pielou’s index (evenness) represents the relative abundance of each taxon in the communities. The influence of age and sex (=cage) was tested on alpha-diversity and beta-diversity metrics using Wilcox univariate nonparametric test and permutational multivariate analysis of variance (PERMANOVA) multivariate test. Age- and sex (=cage)-associated differences in taxa or KO abundance were assessed using the centered log-ratio (CLR) by analysis of composition of microbiomes (ANCOM).

### Mouse treatments

#### In vivo mouse bacterial colonization

10-12-week old, C57BL/6 male mice were administered an antibiotic cocktail (containing 0.5 mg/ml Ampicillin, 0.5 mg/ml Neomycin, 0.25 mg/ml Vancomycin, and 0.5 mg/ml Metronidazole) in drinking water for a total of 2 weeks. After 2 days, mice were administered 0.2 mL of *Clostridium* sp. ASF356 bacterial suspension by oral gavage inside a biological safety cabinet. For robust colonization, oral gavages were repeated for 3 consecutive days, followed by every 7 days for 4 weeks. Mice were maintained on a sterilized standard chow diet (containing phenylalanine 4.15 g/100 g; Research Diets, USA) until they were sacrificed for blood collection and tissue harvesting. To verify the successful colonization, DNA was isolated from fecal content obtained from the distal colon and used for a PCR reaction.

#### In vivo PAA administration

Wild-type or atherosclerosis-prone *Ldlr*-KO mice were administered PAA (50 mg/kg/d, Sigma-Aldrich, P16621) via *i.p.* injection daily for 4 or 8 weeks. Mice were sacrificed for blood collection and harvesting of aortas and PVAT tissues.

#### In vivo senolytic therapy of colonized mice

Antibiotic-pretreated mice were colonized with *Clostridium sp. ASF356* for 4 weeks, followed by treatment with a senolytic cocktail consisting of Dasatinib (5 mg/kg, Sigma-Aldrich, CDS023389) and Quercetin (50 mg/kg, Sigma-Aldrich, Q4951) by oral gavage daily for 3 consecutive days, followed by a 7-day resting period. Mice were maintained on a sterilized standard chow diet (containing phenylalanine 4.15 g/100 g; Research Diets, USA) until they were sacrificed for blood collection and harvesting of aortas and PVAT tissues.

### Cell lines

HAECs (Lonza, CC-2535) were cultured in EBM-2 medium (Lonza, 00190860) supplemented with EGM-2 (10% FBS, 2 mM L-glutamine, 100 μg/mL penicillin-streptomycin, Lonza, CC-3162/6) at 37°C, 5% CO_2_. Cells at passages 4–6 were used as proliferating controls, whereas cells at passages 15–17 represented the replicative senescence state. Cellular senescence was confirmed by proliferative arrest (reduced cell count), SA-β-gal staining, and p21^WAF1/Cip1^ immunoblots, as previously described^2,50^.

Human adipocyte-derived stem cells (hADSC; Lonza, LZ-PT-5006) were cultured in RPMI 1640 medium (Gibco, ThermoFisher Scientific, 11875093) supplemented with 10% FBS at 37°C, 5% CO_2_. Cells were incubated with Preadipocyte Growth Medium (PGM, PromoCell, C-27417) supplemented wiuth SupplementMix (5% FCS, 0.4% ECGF, 10 ng/ml EGF, 1 μg/ml Hydrocortisone, 90 μg/ml Heparin) for 3 days for differentiation to pre-adipocytes. Cells were grown to 90% confluence and incubated with Preadipocyte Differentiation Medium (PDM, PromoCell, C-27437) supplemented with SupplementMix (8 μg/ml d-Biotin, 0.5 μg/ml recombinant human insulin, 400 ng/ml Dexamethasone, 44 μg/ml IBMX, 9 ng/ml L-Thyroxine, 3 μg/ml Ciglitazone) for 6 days. At day 9, adipocytes were treated with Adipocyte Nutrition Medium (ANM, PromoCell, C-39439) and kept on culture for long-term maturation until day 45 (Supplementary Fig. 1a). The maturation was confirmed by qPCR proof for *Ucp1* and *AdipoQ* transcription (Supplementary Fig. 1b). The human adipocyte model captures key metabolic and thermogenic features relevant to PVAT biology.

In Notch signaling experiments, hADSC-derived adipocytes were incubated with 10□μM DAPT ((2*S*)*-N*-[(3,5-Difluorophenyl)acetyl]-L-alanyl-2-phenyl]glycine 1,1-dimethylethyl ester; Tocris Bioscience, 2634), 100 ng/mL human recombinant IL-6 (hrIL6; STEMCELL Technologies, 78050), or 100 μg/mL Tocilizumab (Roche). For insulin signaling studies, cells were incubated in serum-free medium for 3□h, followed by stimulation with 10□nM insulin for 15 min.

### Cellular senescence models

#### H_2_O_2_-induced premature senescence model

Proliferating HAECs or hADSC-derived adipocytes at passages 4-5 were cultured in EGM-2 (+10% FBS) or RPMI (+10% FBS), respectively, followed by treatment with exogenous H_2_O_2_ (50 μM) for 4 h. The culture media were then replaced, and cells were maintained for up to 72 h.

#### Replicative senescence model

Proliferating HAECs (p.4-5) were cultured in EGM-2 (5,000 cells/cm²) and passaged every 48 h until the proliferation is arrested and senescence phenotype is confirmed. As previously established, cellular senescence was verified by SA-β-gal staining and marked reduction in cell count (stained with 0.4% Trypan Blue)^2,50^.

### Senescence Associated-**β**-galactosidase (SA-**β**-gal) assay

SA-β-galactosidase staining was performed using Cellular Senescence assay kit (Merck Millipore, KAA002), as previously described^2,50^. Briefly, cells were fixed and stained with x-gal solution overnight at 37°C. Senescent cells, illuminated as blue-stained cells, were captured under an Agilent BioTek Lionheart FX automated confocal microscope and analyzed using ImageJ software 1.54j (Fiji).

### Cell viability test

Cultured cells were trypsinized, resuspended in culture medium, and mixed 1:1 with 0.4% Trypan Blue solution, as previously described^2,50^. After 3 min incubation at room temperature, viable (unstained) and non-viable (blue-stained) cells were counted under a light microscope. The percentage of viable cells was finally reported.

### Real-time Quantitative PCR

Total RNA from cells and tissues was extracted using TRIzol Reagent^®^ (Sigma-Aldrich, T9424), as previously described^2,50–53^. Quantitative PCR was performed using the Power SYBR Green PCR Master Mix (Thermo Fisher Scientific, 4367659) on a Quant Studio 5 and 7 cyclers system (Life Technologies, ThermoFischer Scientific, Switzerland) according to the manufacturer’s instructions. The sequences of all primers used for qPCR are listed in Supplementary Table 5.

### siRNA transfection

Cells were grown to 70–80% confluence for transfection with specific commercially available ON-TARGETplus HES1 siRNA (SMARTpool 5 nmol, L-007770-02-0005, Dharmacon, Horizon Discovery) using Lipofectamine RNAiMAX transfection reagent (Invitrogen, 13778150). An ON-TARGETplus non-targeting control siRNA (SMARTpool 5 nmol, D-001810-10-05, Dharmacon, Horizon Discovery) was also used as negative control (Supplementary Table 6). The silencing efficiency was confirmed by immunoblots.

### Immunoblotting

Cells and tissues were lysed in ice-cold RIPA buffer (ThermoFisher Scientific, 89901) or nuclear and cytoplasmic extraction reagents (ThermoFisher Scientific, 78835) supplemented with Halt protease and phosphatase inhibitors cocktail (ThermoFisher Scientific, 78442). Total protein was quantitated using the BCA protein assay kit (ThermoFisher Scientific, 23225). Equal amounts of proteins (15-20 µg) were separated on pre-made gels (8-12%, BoltTM Bis-Tris Plus mini protein gels 1.0 mm, ThermoFischer Sceintific, NW00100BOX) in a SDS-PAGE gel electrophoresis system (mini blot module, ThermoFischer Scientific, B1000), followed by electroteranfer and further incubation with specific antibodies. The protein bands were visualized using Amersham Imager 600 (GE Healthcare) and quantitative densitometric analyses were performed using ImageJ software 1.54j (Fiji).

All antibodies used in immunoblotting are listed in Supplementary Table 7.

### Immunofluorescence assay

Aorta and PVAT tissues were harvested from young and old wild-type, *Ldlr*-KO, and p16-3MR mice, fixed in Bouin’s fixative (Muto Pure Chemicals) for 2□h at room temperature^54^. Tissues were processed by paraffin-embedding, followed by sectioning at a thickness of 5-μm. The sections were deparaffinized, unmasked, and blocked with Streptavidin/Biotin blocking solution (Vector Laboratories) and 5% goat serum. The slides were stained with CD31, perilipin, p16^INK4a^, IL-6, or NOTCH1 primary antibodies for 1 h at room temperature, followed by washing with 1x PBST for 3 times. The slides were further incubated with the respective secondary Fluor chrome antibodies (conjugated with AlexaFluor 488 or 594; 1:200) for 30 min at room temperature. Slides were washed with 1x PBST for 3 times, and incubated with mounting media containing DAPI (for nuclear counterstaining) for scanning under fluorescent confocal microscope (Leica TCS-SP8). Another subset of tissue sections were also prepared for hematoxylin and eosin (H&E) staining.

### Aortic root histology

Aortas were collected from wild-type and *Ldlr^−/−^* mice fed a chow diet, Western diet, or PAA. For aortic root histological analysis, tissues were fixed in 4% paraformaldehyde (Santa Cruz Biotechnology, sc-281692) overnight, paraffin-embedded, and sectioned at 5□µm thickness. Sequential sections were stained with H&E for plaque area analysis. The sections were dehydrated through graded ethanol and cleared in xylene. Images were acquired using a Zeiss confocal microscope using the Zen software and analyzed with ImageJ software 1.54j (Fiji). Plaque area was measured on H&E-stained sections.

### Proteome profiler array

Plasma samples were used for profiling inflammatory cytokines and chemokines using a proteome profiler mouse XL cytokine array kit (R&D Systems, ARY028), according to manufacturer instructions^47^. Membranes were visualized using an Amersham Imager 600 (GE Healthcare) and analyzed using ImageJ software 1.54j (Fiji). Intensities were normalized and expressed as relative pixel values.

### IL-6 measurement

Cell culture supernatants were assessed for IL-6 using a Quantikine™ ELISA Human IL-6 Immunoassay kit (R&D Systems, D6050), according to the manufacturer’s instructions. Briefly, Assay Diluent RD1W was added to each well of the 96-well microplate pre-coated with a monoclonal antibody specific for human IL-6. Then, 100□µL of culture supernatant was added and incubated for 2 h at room temperature. Following 2 h incubation with Horseradish Peroxidase (HRP)-conjugated anti-IL-6, substrate solution was added and incubated for 30 min in the dark. Optical density was measured at 450□nm with wavelength correction at 540□nm. IL-6 concentrations (pg/mL) were calculated according to a standard curve.

### Biochemical analysis in the plasma and serum

Total cholesterol and triglycerides were quantified in plasma by colorimetric enzymatic technique (ABX Pentra, HORIBA Medical). Serum insulin was also measured using an enzymatic assay (Crystal Chem, 90080) according to the manufacturer’s protocols.

### Creatinine and BUN measurement

Serum creatinine and BUN were measured using colorimetric detection kits (Cayman Chemical, 700460 and Invitrogen, EIABUN, respectively), according to manufacturers’ instructions. Absorbance was read at 495 nm and 450 nm for creatinine and BUN, respectively, with a Tecan microplate reader.

### Statistics and reproducibility

No statistical methods were used to pre-determine sample sizes, but similar sample sizes were used in previous publications^2,47,50,51^. In TwinsUK cohort, PAA levels were measured in plasma samples of 2,953 subjects by Metabolon using a non-targeted UPLC–MS/MS platform. For the human ASCVD and validation cohorts, no sample size calculation was conducted, as samples were retrospectively collected from observational studies. The cohorts were assembled with the goal of maximizing biological and phenotypic depth rather than targeting a predefined effect size. Categorical data are presented as counts and percentages. PAA concentrations were stratified into tertiles based on their baseline concentration levels. Continuous variables were checked for normality and compared using one-way ANOVA for normally distributed data or the Kruskal-Wallis test for non-normally distributed data. Categorical variables were compared using the chi-square test, Fisher’s Exact test, or Welch’s t-test when expected cell counts were low. To assess a potential association of PAA and traditional risk factors, linear regression either unadjusted or adjusted on a set of confounders (age, sex, smoking, alcohol, family history of CVD, LDL-C, triglycerides, HDL-C, Hb1Ac, CRP, troponin T, and NT-proBNP) was used.

No method of randomization was used to assign animals to experimental groups. No animals or data points were excluded from experiments or analyses. Animals were monitored daily for sickness symptoms and body weight during their lifespan. They were euthanized immediately at the clinical end point when recommended by veterinary and biological service staff members. The investigators in this study were blinded to the conditions of the experiments, except for oral gavage (for *Clos*, PAA, or D+Q administration) to avoid cross-contamination within the groups. *n* represents the number of samples or animals used. At least three independent triplicated experiments were performed for each experimental set-up. Data are expressed as mean ± SD or SEM, and *P*<0.05 was defined as statistically significant. Continuous data were shown as median ± SD or interquartile ranges (IQR) if skewed.

Statistical analysis was performed using GraphPad Prism 9 (v.9.5.1, La Jolla, CA, USA), R (v4.5.1, v4.5.2, 2025), and R package (v3.1.3). Comparisons between groups were conducted using unpaired two-tailed Student’s *t*-test, two-tailed Mann-Whitney *U*-test, two-sided Fisher’s Exact test, two-sided Welch’s t-test, chi-square test, and one-way or two-way ANOVA with Tukey’s post *hoc* test. Heatmaps were generated using the vegan v2.5-5 package (https://Github.com/vegan). Pearson and Spearman rank correlations were used to analyze associations between plasma PAA levels and either age or microbial taxa, respectively. Multiple group comparisons were tested using Kruskal-Wallis and PERMANOVA was used for analysis of alpha- and beta-diversity. Multiple comparisons and associated *P*-values were FDR corrected. Data distribution was assumed to be normal but this was not formally tested.

## Data availability

Source Data and Supplementary Information are provided with this paper. All data from shotgun metagenomics analyses in this study have been deposited in the National Center for Biotechnology Information Sequence Read Archive (SRA) under accession number <u>PRJNA1242241</u>. Screening of K00169 and K00179 was conducted using KEGG annotations, which are accessible at https://ezmeta.unige.ch/CMMG/functional_annotations/mouse/annotations.tsv. The targeted metabolomics data, including Thermo’s raw data files and processed datasets analyzed by Tracefinder 3.2, have been securely stored on an institutional research repository at University of Lausanne. These data are available upon request from the corresponding authors and subject to institutional data-sharing policies. The human study data used in this research are maintained by the Departments of Cardiology and Cardiac Surgery at University Hospital Zurich and Department of Twin Research at King’s College London and are not publicly available due to patient privacy and data protection regulations. Researchers seeking access must apply through the established procedures outlined by the Aging Heart Zurich Cohort (https://www.usz.ch/studie/aging-heart-zuerich-beobachtungsstudie-zur-lebensqualitaet-von-ueber-80jaehrigen-patienten-mit-einer-herzerkrankung/) and the Wellcome Trust, following the guidelines provided at https://twinsuk.ac.uk/resources-for-researchers/access-our-data/. Access is granted upon approval and may require specific ethical and governance clearances. All other data and reagents that support the findings of this study are available upon reasonable request from the corresponding author.

## Acknowledgements

We appreciate Prof. Michael Wannemuehler (Iowa State University) for providing *Clostridium* sp. ASF356.

The authors acknowledge funding from the Swiss National Science Foundation Spark grant (#CRSK-3_229134), the Iten Kohaut Foundation in collaboration with the USZ Foundation (#2512), Novartis Foundation for Medical-Biological Research (#21A053), and the SwissLife Jubiläumsstiftung (#1565) (to S.S.S.S.). K.S. has received support from SwissLife Jubiläumsstiftung (#1438 and #1563) and Wael Almahmeed International Atherosclerosis Society (IAS) Research Fellowship. T.S. is funded by the SwissLife Jubiläumsstiftung (#1564) and Wael Almahmeed International Atherosclerosis Society (IAS) Research Fellowship.

The Department of Twin Research receives support from grants from the Wellcome Trust (212904/Z/18/Z), the Wellcome Leap Dynamic Resilience program (co-funded by Temasek Trust), the Medical Research Council/British Heart Foundation (MR/M016560/1), European Union, Chronic Disease Research Foundation, Zoe Global, Ltd., the National Institutes of Health and Research Clinical Research Facility and Biomedical Research Centre (based at Guy’s and St Thomas’ National Health Service Foundation Trust in partnership with King’s College London). C.M. is supported by the CDRF, y the Italian Ministry of Health–Bando Ricerca Corrente 2023 and by the UK Research Innovation/Medical Research Council (MR/W026813/1 and MR/Y010175).

## Contributions

S.S.S.S. conceptualized and designed the study, and drafted, wrote, and revised the manuscript. K.S., T.S., A.M., and M.S. performed the experiments, including mouse studies, *in vivo* bacterial colonization, senescence and signal transduction studies, senolytic therapy, immunofluorescence assay, and hADSC-adipocyte studies. F.C. and B.P. analyzed shotgun metagenomic data and performed *in vitro* bacterial studies. A.T. and S.L. performed LC-MS/MS quantification of PAA in plasma and culture media. C.M. and X.Z. analyzed data from the human TwinsUK cohort. M.C. and A.A. provided and analyzed p16-3MR mice. L.D.D. and G.D.N. conducted *Ldlr*-KO athero-prone mouse modeling and provided tissues and data. H.R. and O.D. collected aorta and PVAT samples during CABG surgery. S.S.S.S. and M.H. contributed to management and sample collection within Aging Heart Zurich Cohort. S.S.S.S., F.P., and F.R. contributed to interpretation of data. All authors provided intellectual contributions and approved the final manuscript. S.S.S.S. directed and supervised all aspects of the study.

## Corresponding author

Correspondence to Seyed Soheil Saeedi Saravi.

---

**Extended Data Fig. 1. Phenylalanine derivatives are associated with *Clostridium* genus in aged humans. a,** Correlation between plasma PAGln concentrations and chronological age in the TwinsUK aging cohort (*n* = 2,953; male and female). **b,** Correlation between plasma PAGln concentrations and relative abundance of *Clostridium* genus (*n*□=□936, TwinsUK; male and female). **c,** Heatmap shows correlations between plasma PAA or PAGln concentrations and gut bacteria belonging to *Clostridium* genus in human participants (*n*□=□936, TwinsUK; male and female). Statistical analysis was performed using two-tailed Pearson correlation analysis (**a,c**) and linear mixed model-centered log ratio (CLR) transformation (**b**).

**Extended Data Fig. 2. Gut microbiota composition alters with age.** Plots demonstrating age-associated alterations in the abundance of PPFOR-expressing gut microbiota profiles at strain level in in the mouse aging cohort (C57BL/6J male and female; *n* = 10-12). Statistical analysis was performed using analysis of composition of microbiomes (ANCOM) method for microbial abundance analysis. Data are shown as median with min–max; each box plot represents interquartile range (IQR); center lines indicate the median; whiskers extend from the min to max values.

**Extended Data Fig. 3. Aging is associated with aortic PVAT senescence. a-c,** Representative immunoblots and quantification of intensities for the senescence hallmarks CDK inhibitor p16^INK4A^ (**a**) and DNA damage marker γ-H2A.X (**b**), alongside the adipocyte thermogenic marker UCP1 and anti-inflammatory adiponectin (**c**) in tPVAT from aged and young mice (*n* = 6). Error bars represent SD (**a-c**). *P* values were calculated using a two-tailed unpaired Student’s *t*-test (**a-c**).

**Extended Data Fig. 4. PAA induces premature senescence in endothelial cells. a,** Proliferating ECs (passage 5) were treated with PAA (10 μM) or vehicle for 72 h (*n*□=□6 biologically independent samples). **b,c,** Representative bright-field images depicting senescence phenotype (enlarged, flattened, and multinucleated) (**b**) and SA-β-gal^+^ cells, confirming senescence-associated lysosomal alterations (**c**). **d,** Immunostaining for the proliferation marker Ki67. **e,** qPCR demonstrates transcriptional alterations of the SASP components *IL1A*, *IL1B*, and *IL6* (*n* = 6). Data were determined in 15-18 micrographs and represent triplicated biologically independent experiments (**b-d**). Scale bar, 50, 100, and 400 μm (**b-d**). Error bars represent SD (**c,e**). *P* values were calculated using a two-tailed unpaired Student’s *t*-test (**c,e**). Image created with https://BioRender.com (**a**).

**Extended Data Fig. 5. Senolytics attenuate PAA-triggered adipocyte senescence. a,** hADSC-derived adipocytes were treated with the conditioned medium (CM) derived from ECs (passage 5) exposed to PAA (10 μM, PAA-CM) or vehicle (control-CM) for 72 h in the presence of Dasatinib+Quercetin (D□+□Q: 500□nM + 20□μM) for 24 h. Cells were then subjected to senescence hallmarks profiling. **b,** Cell counts demonstrate the viability percentage (%) in PAA-CM- or control-CM-exposed adipocytes in response to D+Q (*n*□=□6). All experiments were triplicated independently. *P* values were calculated using a one-way ANOVA followed by Tukey’s post *hoc* test (**b**). Image created with https://BioRender.com (**a**).

**Extended Data Fig. 6. PAA-induced EC senescence promotes preadipocyte senescence. a,** hADSC-derived preadipocytes were treated with the conditioned medium (CM) derived from ECs (passage 5) exposed to PAA (10 μM, PAA-CM, for 72 h) or H_2_O_2_ (50 μM, H_2_O_2_-CM, for 4 h). **b,c,** Bar charts represent transcriptional alterations of the CDK inhibitors *Cdkn2a*, *Cdkn2d*, and *Cdkn1a* (**b**), as well as, the SASP components *Il1b*, *Il6*, and *Ccl2* (**c**), as assessed by qPCR (*n* = 6). **d,** Representative bright-field images demonstrating senescence phenotype (upper) and SA-β-gal^+^ cells (lower). Data were determined in 15-18 micrographs and represent triplicated biologically independent experiments (**b-d**). Scale bars, 200 and 400 μm (**d**). Error bars represent SD (**b-d**). *P* values were calculated using a two-tailed unpaired Student’s *t*-test (**b,c**) and a one-way ANOVA followed by Tukey’s post *hoc* test (**d**). Image created with https://BioRender.com (**a**).

**Extended Data Fig. 7. hrIL6 promotes adipocyte senescence and dysfunction through Notch1 activation. a,** hADSC-derived adipocytes were pretreated with the NOTCH inhibitor, DAPT (10 μM), or vehicle for 72 h, followed by treatment with hrIL6 (100 ng/mL) for 2 h. **b,** Representative immunoblots and quantification of intensities for NOTCH1 (*n* = 6). **c,** Representative bright-field images show the number of SA-β-gal^+^ cells (%) (*n* = 6). **d,** Immunobloting for the insulin signaling pathway (*n* = 6). Data were determined in 15-18 micrographs and represent triplicated biologically independent experiments (**b-d**). Scale bar, 200 μm (**c**). Error bars represent SD (**b-d**). *P* values were calculated using a two-tailed unpaired Student’s *t*-test (**b-d**). Image created with https://BioRender.com (**a**).

**Extended Data Fig. 8. Senolytics improve systemic metabolic health in *Clos*-colonized mice. a,** Body weight (g) at baseline (pre) and after treatment (post). Young mice were colonized with *Clostridium* sp. ASF356 or vehicle for 4 weeks with or without administration of senolytic D+Q cocktail (5 + 50 mg/kg/d, for 3 days) (*n*□=□5-6). **b,** Serum insulin levels in these mice (*n* = 5-6). Error bars represent SD (**a,b**). *P* values were calculated using a one-way ANOVA followed by Tukey’s post *hoc* test (**a,b**).

**Extended Data Fig. 9. Clearance of senescent cells perturbs the pro-atherogenic microenvironment. a,** p16-3MR mice were administered with GCV (25 mg/kg) or vehicle (PBS) via daily *i.p.* injections for 5 days. **b,** H&E staining (left) and immunostaining (right) of aortic and tPVAT tissues (*n*□=□5). **c,** Representative bright-field images of SA-β-gal^+^ cells. **d,** qPCR demonstrates transcriptional alterations of pro-atherogenic markers *Il1b*, *Il6*, *Tnfa*, *Vcam1*, *Mmp2*, and *Mmp9* in aortas (left) and thermogenic and inflammatory genes *Ucp1*, *AdipoQ*, *Leptin*, *Il1b*, *Il6*, *Ccl2*, and *Tnfa* in tPVAT (right) (*n* = 5). Data were determined in 15-18 micrographs and represent triplicated biologically independent experiments (**b-d**). Scale bar, 200 μm. Error bars represent SD (**c,d**). *P* values were calculated using a two-tailed unpaired Student’s *t*-test (**c,d**). Image created with https://BioRender.com (**a**).

**Extended Data Fig. 10. Systemic metabolic impairment in atherosclerosis-prone *Ldlr^-/-^*mice. a-c,** Serum insulin levels (**a**), body weight (**b**), and plasma cholesterol and triglycerides (**c**) in *Ldlr^−/−^* and WT mice were fed chow (for 8 weeks) or Western diet (for 12 weeks). WC: WT + chow diet; WW: WT + Western diet; LC: *Ldlr^−/−^*+ chow diet; LW: *Ldlr^−/−^* + Western diet. Error bars represent SD (**a-c**). *P* values were calculated using a one-way ANOVA followed by Tukey’s post *hoc* test (**a-c**).

**Extended Data Fig. 11. PAA triggers vascular senescence, driving atherosclerosis in *Ldlr^-/-^* mice.** H&E staining (left) and immunostaining (right) of aortas from chow-fed *Ldlr^−/−^* mice treated with PAA or vehicle for 8 weeks (*n*□=□5). Arrowheads indicate plaque areas. Data were determined in 8-10 micrographs. Scale bars, 20 and 200 μm.

## References

1. Ding, Y., et al. Comprehensive human proteome profiles across a 50-year lifespan reveal aging trajectories and signatures. Cell. 188, 5763–5784.e26 (2025).

2. Saeedi Saravi, S.S., et al. Gut microbiota-dependent increase in phenylacetic acid induces endothelial cell senescence during aging. Nat Aging. 5, 1025–1045 (2025).

3. Ruschitzka, F., et al. Gut Microbial metabolites connection to cardiovascular disease call a gusty therapeutic apparoches. J Clin Invest. 135, e201468 (2025).

4. Saeedi Saravi, S.S. Metabolites matter for gut microbiota as a modifiable risk factor in cardiovascular diseases. Nat Rev Cardiol 22, 610 (2025).

5. Mastrangelo, A., et al. Imidazole propionate is a driver and therapeutic target in atherosclerosis. Nature. 645, 254–261 (2025).

6. Nemet, I., et al. Atlas of gut microbe-derived products from aromatic amino acids and risk of cardiovascular morbidity and mortality. Eur Heart J. 44, 3085–3096 (2023).

7. Zhu, Y., et al. Two distinct gut microbial pathways contribute to meta-organismal production of phenylacetylglutamine with links to cardiovascular disease. Cell Host Microbe. 31, 18–32.e9 (2023).

8. Yang, H., et al. Gut microbial-derived phenylacetylglutamine accelerates host cellular senescence. Nat Aging. 5, 401–418 (2025).

9. Barinda, A.J., et al. Endothelial progeria induces adipose tissue senescence and impairs insulin sensitivity through senescence associated secretory phenotype. Nat Commun. 11, 481 (2020).

10. Suda, M., et al. Endothelial senescent-cell-specific clearance alleviates metabolic dysfunction in obese mice. Cell Metab. 37, 2455–2465.e6 (2025).

11. Antoniades, C., et al. Perivascular adipose tissue as a source of therapeutic targets and clinical biomarkers: A clinical consensus statement from the European Society of Cardiology Working Group on Coronary Pathophysiology and Micro-circulation. Eur Heart J. 44, 3827–3844 (2023).

12. Akawi, N., et al. Fat-Secreted Ceramides Regulate Vascular Redox State and Influence Outcomes in Patients With Cardiovascular Disease. J Am Coll Cardiol. 77, 2494–2513 (2021).

13. Antonopoulos, A.S., et al. Detecting human coronary inflammation by imaging perivascular fat. Sci Transl Med. 9, eaal2658 (2017).

14. Akoumianakis, I., et al. Perivascular adipose tissue as a regulator of vascular disease pathogenesis: identifying novel therapeutic targets. Br J Pharmacol. 174, 3411–3424 (2017).

15. Nemet, I., et al. A Cardiovascular disease-linked gut microbial metabolite acts via adrenergic receptors. Cell. 180, 862–877 (2020).

16. Boucher, J.M., et al. Pathological Conversion of Mouse Perivascular Adipose Tissue by Notch Activation. Arterioscler Thromb Vasc Biol. 40, 2227–2243 (2020).

17. Potts, C., et al. NOTCH signaling networks in perivascular adipose tissue. Arterioscler Thromb Vasc Biol. 45, 845–856 (2025).

18. Zhang, Y., et al. Inflammatory Cytokine Interleukin-6 (IL-6) promotes the proangiogenic ability of adipose stem cells from obese subjects *via* the IL-6 signaling pathway. Curr Stem Cell Res Ther. 18, 93–104 (2023).

19. Chatterjee, T.K., et al. Proinflammatory phenotype of perivascular adipocytes: influence of high-fat feeding. Circ Res. 104, 541–549 (2009).

20. Xu, C., et al. The role of cellular senescence in cardiovascular disease. Cell Death Discov. 11, 431 (2025).

21. Song, J., et al. Age-associated adipose tissue inflammation promotes monocyte chemotaxis and enhances atherosclerosis. Aging Cell. 22, e13783 (2023).

22. Honda, S., et al. Cellular senescence promotes endothelial activation through epigenetic alteration, and consequently accelerates atherosclerosis. Sci Rep. 11, 14608 (2021).

23. Han, Y. & Kim, S.Y. Endothelial senescence in vascular diseases: current understanding and future opportunities in senotherapeutics. Exp. Mol. Med. 55, 1–12 (2023).

24. Eroglu, E., et al. Discordance between eNOS phosphorylation and activation revealed by multispectral imaging and chemogenetic methods. Proc. Natl. Acad. Sci. USA. 116, 20210–20217 (2019).

25. Koplev, S., et al. A mechanistic framework for cardiometabolic and coronary artery diseases. Nat Cardiovasc Res. 1, 85–100 (2022).

26. Jie, Z., et al. The gut microbiome in atherosclerotic cardiovascular disease. Nat Commun. 8, 845 (2017).

27. Wang, T., et al. Divergent age-associated and metabolism-associated gut microbiome signatures modulate cardiovascular disease risk. Nat Med. 30, 1722–1731 (2024).

28. Chajadine, M., et al. Harnessing intestinal tryptophan catabolism to relieve atherosclerosis in mice. Nat Commun. 15, 6390 (2024).

29. Nageswaran, V., et al. Gut Microbial Metabolite Imidazole Propionate Impairs Endothelial Cell Function and Promotes the Development of Atherosclerosis. Arterioscler Thromb Vasc Biol. 45, 823–839 (2025).

30. Kasahara K, et al. Interactions between *Roseburia intestinalis* and diet modulate atherogenesis in a murine model. Nat Microbiol. 3, 1461–1471 (2018).

31. Tang, W.H.W., et al. Intestinal Microbial Metabolism of Phosphatidylcholine and Cardiovascular Risk. N Engl J Med. 368, 1575–1584 (2013).

32. Yang, C., et al. Notch signaling regulates mouse perivascular adipose tissue function via mitochondrial pathways. Genes. 14, 1964 (2023).

33. Bi, P., et al. Inhibition of Notch signaling promotes browning of white adipose tissue and ameliorates obesity. Nat Med. 20, 911–918 (2014).

34. Jin, Y., et al. PVAT-conditioned media from Dahl S rats on high fat diet promotes inflammatory cytokine secretion by activated T cells prior to the development of hypertension. PLoS One. 19, e0302503 (2024).

35. Saeedi Saravi, S. S. & Feinberg, M. W. Can removal of zombie cells revitalize the aging cardiovascular system? Eur. Heart J. 45, 867–869 (2024).

36. Bloom, S., I., et al. Mechanisms and consequences of endothelial cell senescence. Nat. Rev. Cardiol. 20, 38–51 (2023).

37. Fromentin, S. et al. Microbiome and metabolome features of the cardiometabolic disease spectrum. Nat. Med. 28, 303–314 (2022).

38. Kaasenbrood, L., et al. Distribution of estimated 10-year risk of recurrent vascular events and residual risk in a secondary prevention population. Circulation. 134, 1419–1429 (2016).

39. Nelson, K., et al. Low-dose colchicine for secondary prevention of coronary artery disease. J Am Coll Cardiol. 82, 648–660 (2023).

40. Lloyd-Jones, D.M., et al. 2017 focused update of the 2016 ACC expert consensus decision pathway on the role of non-statin therapies for LDL-cholesterol lowering in the management of atherosclerotic cardiovascular disease risk: a report of the American College of Cardiology Task Force on Expert Consensus Decision Pathways. J Am Coll Cardiol. 70, 1785–1822 (2017).

41. Romano, K.A., et al. Gut microbiota-generated phenylacetylglutamine and heart failure. Circ Heart Fail. 16, e009972 (2023).

42. Tang, W.H.W., et al. Prognostic value of gut microbe-generated metabolite phenylacetylglutamine in patients with heart failure. Eur J Heart Fail. 26, 233–241 (2024).

43. Allemann, S., et al. The gut-microbiome derived phenylacetylglutamine predicts adverse events in patients with acute coronary syndromes. Eur Heart J. 45, ehae666.1600 (2024).

44. https://twinsuk.ac.uk/

45. Keiser, S., et al. Comprehensive mouse microbiota genome catalog reveals major difference to its human counterpart. PLoS Comput Biol. 18, e1009947 (2022).

46. https://www.usz.ch/studie/aging-heart-zuerich-beobachtungsstudie-zur-lebensqualitaet-von-ueber-80jaehrigen-patienten-mit-einer-herzerkrankung/

47. Saeedi Saravi, S. S., et al. Dietary omega-3 fatty acid suppresses age-associated thrombotic potential via gut microbiota modulation. iScience. 24, 102897 (2021).

48. Blanco-Míguez, A., et al. Extending and improving metagenomic taxonomic profiling with uncharacterized species using MetaPhlAn 4. Nat Biotechnol. 41, 1633–1644 (2023).

49. Visconti, A., et al. Interplay between the human gut microbiome and host metabolism. Nat Commun. 10, 4505 (2019).

50. Shabanian, K., et al. AQP1 differentially orchestrates endothelial cell senescence. Redox Biol. 76,103317 (2024).

51. Saeedi Saravi, S.S., et al. Long-term dietary n3 fatty acid prevents aging-related cardiac diastolic and vascular dysfunction. Vasc Pharmacol. 150, 107175 (2023).

52. Spyropoulos, F., et al. Metabolomic and transcriptomic signatures of Chemogenetic heart failure. Am J Physiol Heart Circ Physiol. 322, H451–H465 (2021).

53. Sorrentino, A., et al. Reversal of heart failure in a chemogenetic model of persistent cardiac redox stress. Am J Physiol Heart Circ Physiol. 317, H617–H626 (2019).

54. Kawamoto, S., et al. Bacterial induction of B cell senescence promotes age-related changes in the gut microbiota. Nat Cell Biol. 25, 865–876 (2023).

